# Major orchestration of shikimate, early phenylpropanoid and stilbenoid pathways by Subgroup 2 R2R3-MYBs in grapevine

**DOI:** 10.1101/2020.12.31.424746

**Authors:** Luis Orduña, Miaomiao Li, David Navarro-Payá, Chen Zhang, Antonio Santiago, Pablo Romero, Živa Ramšak, Gabriele Magon, Janine Höll, Patrik Merz, Kristina Gruden, Alessandro Vannozzi, Dario Cantu, Jochen Bogs, Darren C. J. Wong, Shao-shan Carol Huang, José Tomás Matus

## Abstract

The stilbenoid pathway is responsible for the production of resveratrol and its derivatives in grapevine. A few transcription factors (TFs) have been previously identified as regulators of this pathway but the extent of this control is yet to be fully understood. Here we demonstrate how DNA affinity purification sequencing (DAP-Seq) allows for genome-wide TF binding site interrogation in a non-model species. We obtained 5,190 and 4,443 binding events assigned to 4,041 and 3,626 genes for MYB14 and MYB15, respectively (around 40% of peaks being located within -10kb of transcription start sites). DAP-Seq of MYB14 and MYB15 was combined with aggregate gene centred co-expression networks built from more than 1,400 transcriptomic datasets from leaves, fruits and flowers to narrow down bound genes to a set of high confidence targets. The analysis of MYB14, MYB15 and MYB13, a third uncharacterised member of Subgroup 2 (S2), showed that in addition to the few previously known stilbene synthase (*STS* ) targets, these three regulators bind to 30 out of 47 *STS* family genes. Moreover all three MYBs bind to several *PAL*, *C4H* and *4CL* genes, in addition to shikimate pathway genes, the WRKY03 stilbenoid co-regulator and novel resveratrol-modifying gene candidates amongst which *ROMT2* -*3* were validated enzymatically. A high proportion of DAP-Seq bound genes was induced in the activated transcriptomes of transient *MYB15* -overexpressing stilbenoid-producing grapevine leaves, validating our methodological approach for identifying gene regulatory networks of specialised metabolism. Overall, MYB genes from Subgroup 2 appear to play a key role in binding and directly regulating several primary and secondary metabolic steps leading to an increased flux towards stilbenoid production.

## Introduction

The evolved complexity of plant specialised metabolism can be traced back to the colonisation of dry land by green algal-derived ancestors (Waters, 2003). Since their origin, land plants have been accompanied by a persistent range of abiotic and biotic stresses such as UV radiation, desiccation, or unfavourable microbial communities that have led to the selective emergence of novel protective metabolites derived from primary metabolism (Kenrick and Crane, 1997). A clear example of this is the phenylpropanoid pathway, which is responsible for synthesising a plethora of vital compounds such as lignans and flavonoids, all of which start their biosynthesis from the catabolism of phenylalanine and tyrosine. This pathway is a major source of aromatic secondary metabolites in plants, with many of their roles related to tolerance and adaptation to the above-mentioned stresses (e.g. anthocyanins protecting against excessive radiation). Whilst some branches of the pathway are ubiquitous in the plant kingdom (such as those producing flavonoids), others such as the stilbene pathway are restricted to a small number of species across at least 10 unrelated families including Vitaceae (e.g. *Vitis vinifera* L.) and Moraceae (e.g. *Morus alba*) (Dubrovina and Kiselev, 2017). Grapevine is not only an important crop species but an interesting model to study the complexity of the stilbene pathway given the remarkable expansion of the stilbene synthase (*STS* ) family in its genome through segmental and tandem gene duplications, reaching up to a total of 47 genes (Parage et al., 2012; Vannozzi et al., 2012), albeit 12 of these are considered pseudogenes.

Stilbenes are phytoalexins; small, lipophilic compounds with key roles in plant defence, which accumulate in response to a range of abiotic and biotic stresses. In recent years, the grapevine *STS* gene family has been studied to reveal a high degree of responsiveness and effectiveness against different biotic or abiotic stresses. For instance, ectopic expression of *VqSTS36* from the Chinese wild species *Vitis quinquangularis*, in both *Arabidopsis* and tomato, enhanced resistance to powdery mildew and osmotic stress (Huang et al., 2018), whilst the expression of *VqSTS29* in *Arabidopsis* also led to powdery mildew resistance (Xu et al., 2019). Moreover, induction of *STS* gene expression in different *V. vinifera* tissues has been observed in response to fungal infection, UV-C or heat treatments (Vannozzi et al., 2012; Yin et al., 2016; Lecourieux et al., 2017).

STS enzymes are direct competitors of chalcone synthases (CHSs) for pathway precursors. Both proteins are closely related type-III polyketide synthases, generating a tetraketide intermediate from the condensation of p-coumaroyl-CoA with 3 molecules of malonyl-CoA, which depending on STS/CHS activity, will generate resveratrol or naringenin chalcone, respectively, thus defining the entry point of the stilbene and flavonoid branches. Different stimuli have been shown to favour one branch over the other, e.g. UV-C irradiation and downy mildew infection promote *STS* gene expression whilst downregulating *CHS* expression (Vannozzi et al., 2012).

The main and first stilbene produced in grape tissues is resveratrol, a well-known nutraceutical with many characterised properties ranging from antioxidant to antiviral activities (e.g. it has been recently shown to inhibit SARS-CoV-2 *in vitro* replication in human lung cells (Pasquereau et al., 2021)). Resveratrol is derivatised into a broad range of stilbenes such as pterostilbene, viniferins, piceid, and piceatannol which involve methoxylation, oligomerisation, glucosylation and hydroxylation, respectively. The enzymes catalysing these reactions are mostly uncharacterised in grapevine except for a resveratrol O-methyltransferase (ROMT1) responsible for the production of pterostilbene (Schmidlin et al., 2008) and a resveratrol glucosyl transferase (Rs-GT) in *Vitis labrusca*, leading to the production of piceid (Hall and De Luca, 2007). Hydroxylation of resveratrol into piceatannol could be carried out by cytochrome P450 oxidoreductases but no candidates have yet been identified in grape.

The stilbene pathway in grapevine is mainly regulated at the transcriptional level through transcription factors (TFs). In particular, the R2R3-type MYB14 and MYB15 (members from Subgroup 2) have been shown to specifically activate a few *STS* promoters (*STS29* and *STS41* ) in transient reporter assays (Höll et al., 2013), but modulation of other stilbenoid branch enzymes remains unexplored. Furthermore, gene co-expression networks (GCNs) and further correlation with stilbene accumulation have pointed out MYB13 as an additional putative regulator of stilbene accumulation (Wong et al., 2016) although it has not been yet validated *in planta*. We have initially explored TF regulatory networks interrogated by the use of GCNs, leading to the identification of members of the *AP2* /*ERF*, *bZIP* and *WRKY* gene families as potential regulators of *STS* expression (Wong and Matus, 2017), however, experimental evidence of binding is necessary to prove regulatory causality. Nevertheless, systems biology approaches initially conducted in (Wong et al., 2016), applied to the regulation of transcription in grapevine, have paved the way for the functional characterisation of additional stilbene pathway regulators such as bZIP1, ERF114, MYB35A and WRKY53 in recent years (Wang et al., 2019; Wang and Wang, 2019; Vannozzi et al., 2018), proving their efficacy in hypothesis-driven research for TF discovery. Interestingly, one of the newly identified stilbene regulators, WRKY53, binds to a subset of *STS* genes (*STS32* and *STS41* ) and it is thought to form a regulatory complex with MYB14 and MYB15, probably increasing its activity (Wang et al., 2020). Moreover, WRKY03 has also been shown to work in synergy with MYB14 in the up-regulation of *STS29* expression (Vannozzi et al., 2018).

Amongst the identified *STS* regulators, MYB14 and MYB15 have been proposed as upstream TFs in the regulatory cascade governing stilbenoid accumulation mainly due to their rapid activation response, however, without any experimental validation. The detailed characterisation of these two potential high-hierarchy regulators is hence of great importance. In addition, no other processes controlled by these TFs have been identified. This study combines genome-wide TF binding-site interrogation using DNA affinity purification sequencing (DAP-Seq), and aggregate whole genome co-expression networks to lay out MYB14 and MYB15 cistrome landscapes and identify their complete repertoire of target genes. We have also analysed the yet uncharacterised MYB13. Our results suggest that these three MYBs bind to regulatory elements in most members of the *STS* gene family, other shikimate and early phenylpropanoid genes as well as to WRKY regulators of stilbene synthesis, representing major regulators of this specialised metabolic pathway.

## Results

### *MYB14* and *MYB15* bind proximal upstream regions of a large set of *STS* genes

Our DAP-Seq analysis reported 5,190 and 4,443 TF binding events, i.e. peaks (Fig. 1a, Dataset S1), which were assigned to 4,041 and 3,626 different genes for *MYB14* and *MYB15*, respectively. An initial inspection of all binding events showed that 73% and 75% of the peaks are present between 10 kb upstream of transcription start sites (TSSs) and 2kb downstream of annotated gene ends for *MYB14* and *MYB15*, respectively. A total of 30% and 32% of *MYB14* and *MYB15* peaks, respectively, are found within 5kb upstream of TSSs. *MYB14* and *MYB15* shared a total of 2,709 bound genes as well as an almost identical DNA binding motif obtained from the enrichment analysis of the top 600 most significant peaks sequences.(Fig. 1b). The cistromes of *MYB14* and *MYB15* Arabidopsis orthologues have not been previously studied.

**Figure 1:**
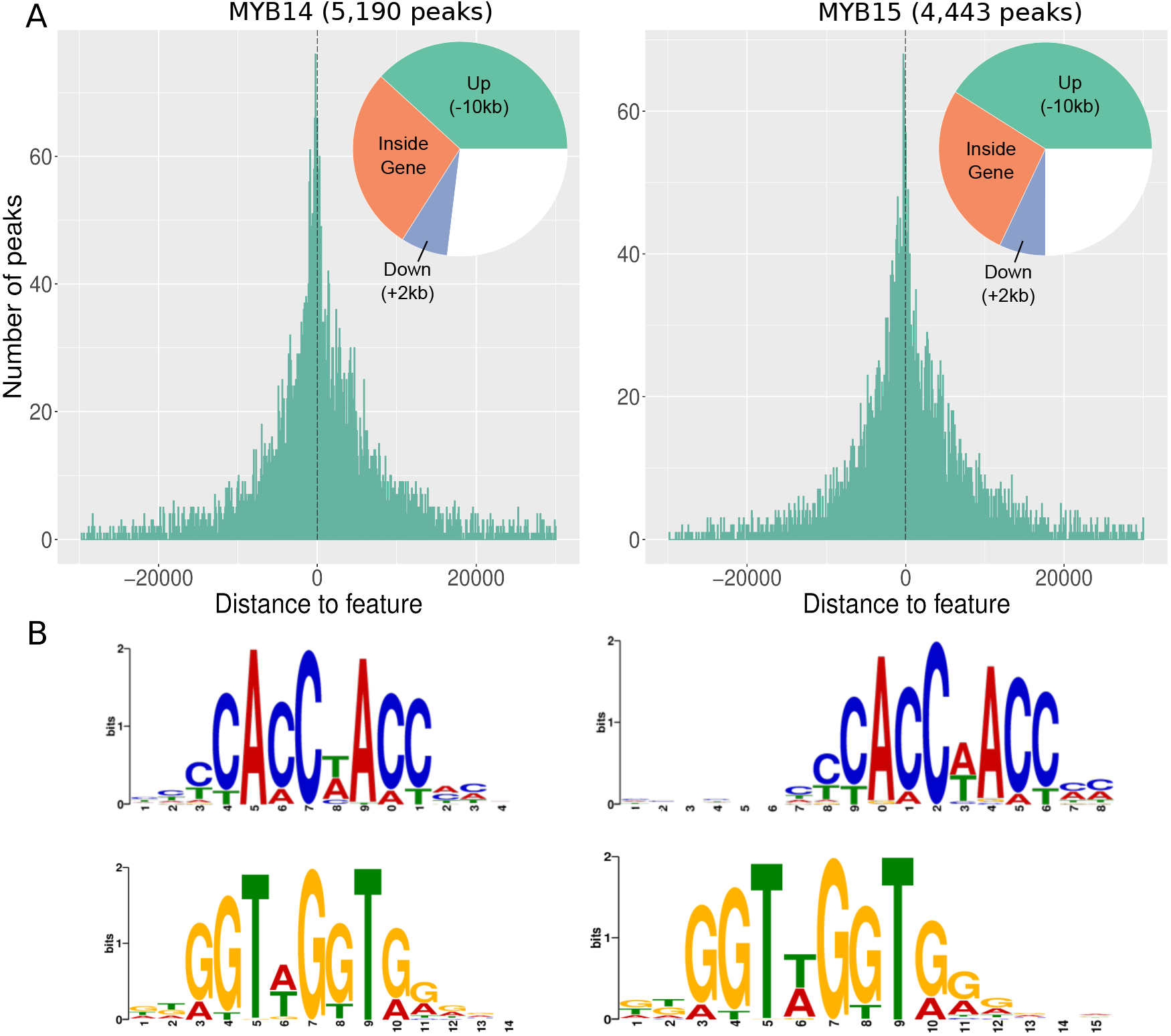
MYB14 and MYB15 DAP-Seq derived cistrome landscapes in *V. vinifera* cultivar (cv.) ‘PN40024’. (a) DNA binding events with respect to all transcription start sites (TSSs) of assigned genes. The proportion of binding peaks 10kb upstream of TSSs, inside genes or 2kb downstream of genes are represented within the pie-charts in green, orange and blue, respectively. (b) *De novo* binding motifs, forward and reverse, obtained from the top 600 scoring peaks of MYB14 and MYB15 using MEME suite.

A closer look at the *STS* gene family revealed that 22 out of the 47 family members have MYB14/MYB15 DAP-Seq peaks associated to them (including 5 out of 13 pseudogenes). Examining the *STS* genomic regions revealed clear DNA binding signals upstream of TSSs for both MYB14 and MYB15 TFs which was not observed in the pIX-HALO non-specific DNA binding control (Fig. 2). A selection of housekeeping genes was used as a negative control with no specific DNA binding of MYB14 or MYB15 TFs around their TSSs. Interestingly, many DAP-Seq bound genes belong to the shikimate pathway or even to the first committed steps of the phenylpropanoid pathway. In addition to *STSs* other enzyme categories such as O-methyltransferases and glucosyltransferases are also represented amongst DAP-Seq bound genes.

**Figure 2:**
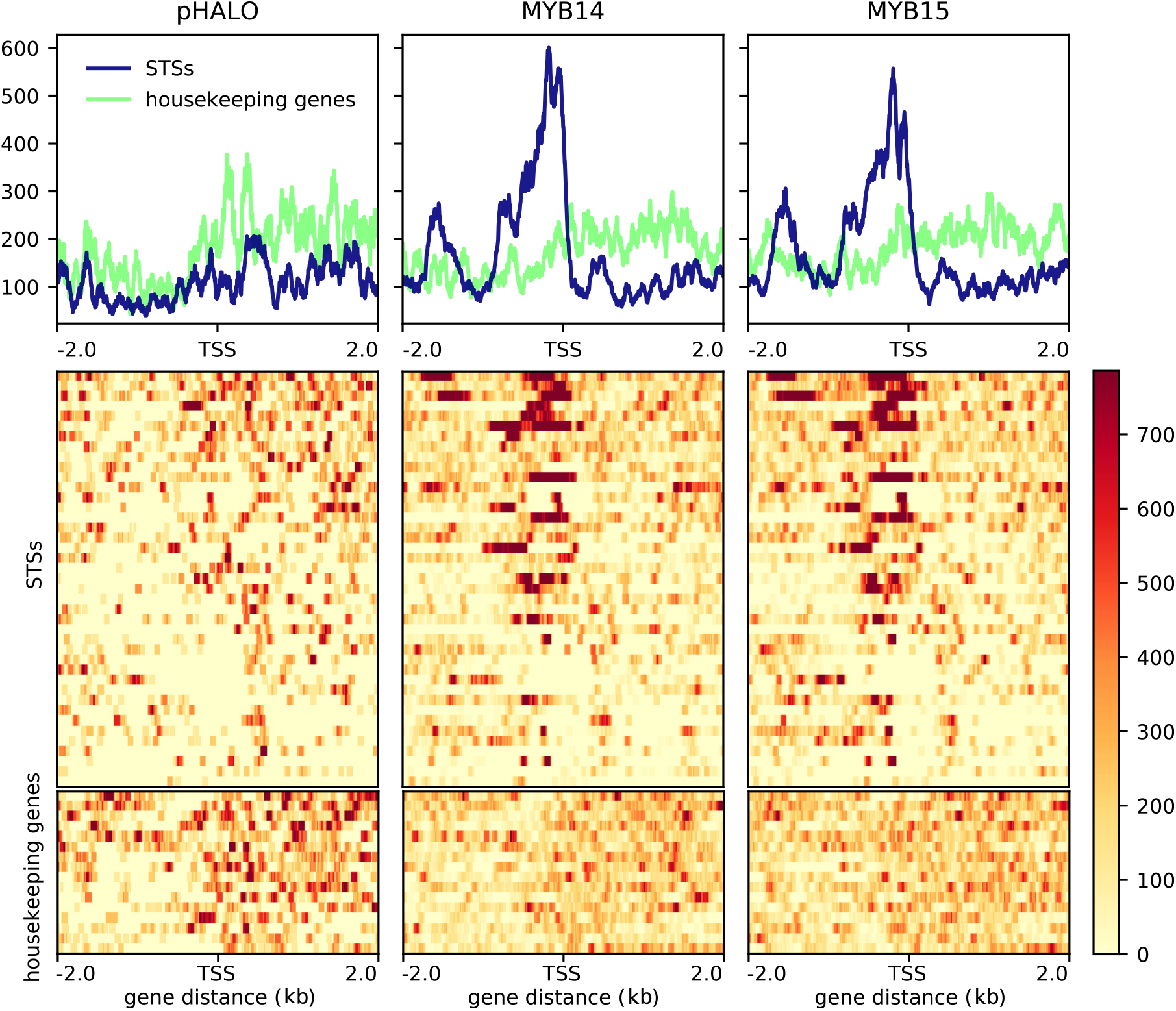
MYB14 and MYB15 DNA binding events in promoter regions of *STS* genes. DAP-Seq binding signal for MYB14 and MYB15 at -2kb and +2kb from the TSS of *STS* genes is high when compared to background housekeeping genes both in the density plots, which show the average binding signal, and the heatmaps showing individual gene profiles. Housekeeping genes are listed in Dataset S1. The figures were generated using the Deeptools suite v.3.3.2, computing and normalizing the coverage for each BAM file with a BinSize = 10 and RPKM normalisation. RPKM value for each bin is the average between the 10 positions that define each bin. BigWig files for the individual replicates for each TF were merged using bigWigMerge v.2 and bedGraphToBigWig v.4.

### Inspection of *MYB14* and *MYB15* gene-centred co-expression networks

We generated condition-dependent leaf and fruit aggregate whole genome co-expression networks using public RNA-Seq data uploaded in SRA, composed of more than 1,400 runs belonging to more than 70 independent experiments (Dataset S2). Network performance was assessed through the area under the receiver operator characteristic curve (AUROC) measurement of the final aggregate network, obtaining AUROC values of `≈` 0.75. The final fruit/flower network consisted of 31,723 genes as individual nodes and 13,323,660 co-expression connections as directed edges (weighted by frequency of co-expression across the selected SRA studies). The leaf network consisted of 31,759 genes and 13,338,780 edges. The two networks share a total of 1,795,970 common interactions, showing a great number of tissue-specific co-expression connections. Gene co-expression networks (GCNs) centred on individual genes were extracted from the final aggregate leaf and fruit/flower networks in the form of the top 420 interactions for each particular gene (corresponding to TOP1% of the total grapevine genes according to the V.Cost annotation). Since *MYB14* and *MYB15* have very low expression in flower tissues, any detected co-expression relationship with other genes in the whole genome network are most surely attributable to fruit samples, therefore, fruit/flower GCNs are referred from here on as fruit GCNs.

Individual *MYB14* and *MYB15* gene centred fruit GCNs revealed that both TFs shared 146 genes out of the 420 GCN members. In addition, *MYB14* is present in the GCN of *MYB15* and *vice versa*. We found 13 *STS* genes present in both fruit GCNs. In addition, 28 *STS* genes are exclusively present in the fruit GCN of *MYB14* meaning that almost all known *STS* genes in the grapevine genome (41 out of 47) are present in any one of the fruit GCNs. The six *STS* genes which are not present in either TF-centred GCN are *STS4* /*11* /*33* /*34* /*44*, all of which are putative pseudogenes. Interestingly, *MYB15* does not have any exclusive *STSs* in its fruit GCN, suggesting a more direct relationship of *MYB14* with this gene family. Other notable genes which are common to both GCNs are *PALs*, *WRKYs* and other genes related to secondary metabolism. There are five *PALs* shared by both fruit GCNs, whilst the *MYB14* and *MYB15* fruit GCNs have two and one exclusive *PAL* gene, respectively. A comparison of *MYB14* and *MYB15* leaf GCNs revealed 180 common genes out of 420, with *MYB14* and *MYB15* again being present in each other’s GCNs. Regarding *STS* genes, 28 out of 47 are present in both MYB GCNs, while 4 and 8 are exclusive of *MYB14* and *MYB15*, respectively. The presence of other notable genes of the secondary metabolism pathways in both GCNs, such as *PALs* and *WRKYs*, is also remarkable.

Results in both leaf and fruit GCNs greatly overlap with DAP-Seq data, providing transcriptional regulation evidence for many co-expression relationships (Fig. 3). The total number of co-expressed genes in both fruit GCNs is of 694 out of which 23% were identified also by DAP-Seq. In the case of the specific *MYB14* and *MYB15* fruit GCNs, 24% and 23% of genes are DAP-Seq bound. A similar tendency is observed for the comparison between leaf GCNs and DAP-seq data. Moreover, amongst these are genes of interest coding for TFs such as WRKYs, NACs or enzymes within the shikimate, early phenylpropanoid or stilbenoid pathways. Most of the genes identified both by DAP-Seq and co-expression analyses had the MYB binding motifs identified within 5kb upstream of TSSs and belonged to shikimate, early phenylpropanoid and stilbenoid pathways. An overlap of this data with the grapevine reference gene catalogue v1.1 available at Integrape (http://www.integrape.eu/index.php/resources/genomes/) revealed a considerable representation of secondary metabolism genes. All the provided gene symbols in this study are in accordance with the reference gene catalogue (v1.1). Bearing in mind that *MYB14* and *MYB15* may be major regulators and hence be co-expressed with a large number of genes, a fraction of biologically relevant co-expressed genes is expected to be absent in their GCNs, which have a 420 gene cut-off. Therefore, to further predict a reliable list of targets, the GCNs of DAP-Seq bound genes were also inspected to interrogate the presence of either *MYB14* or *MYB15*.

**Figure 3:**
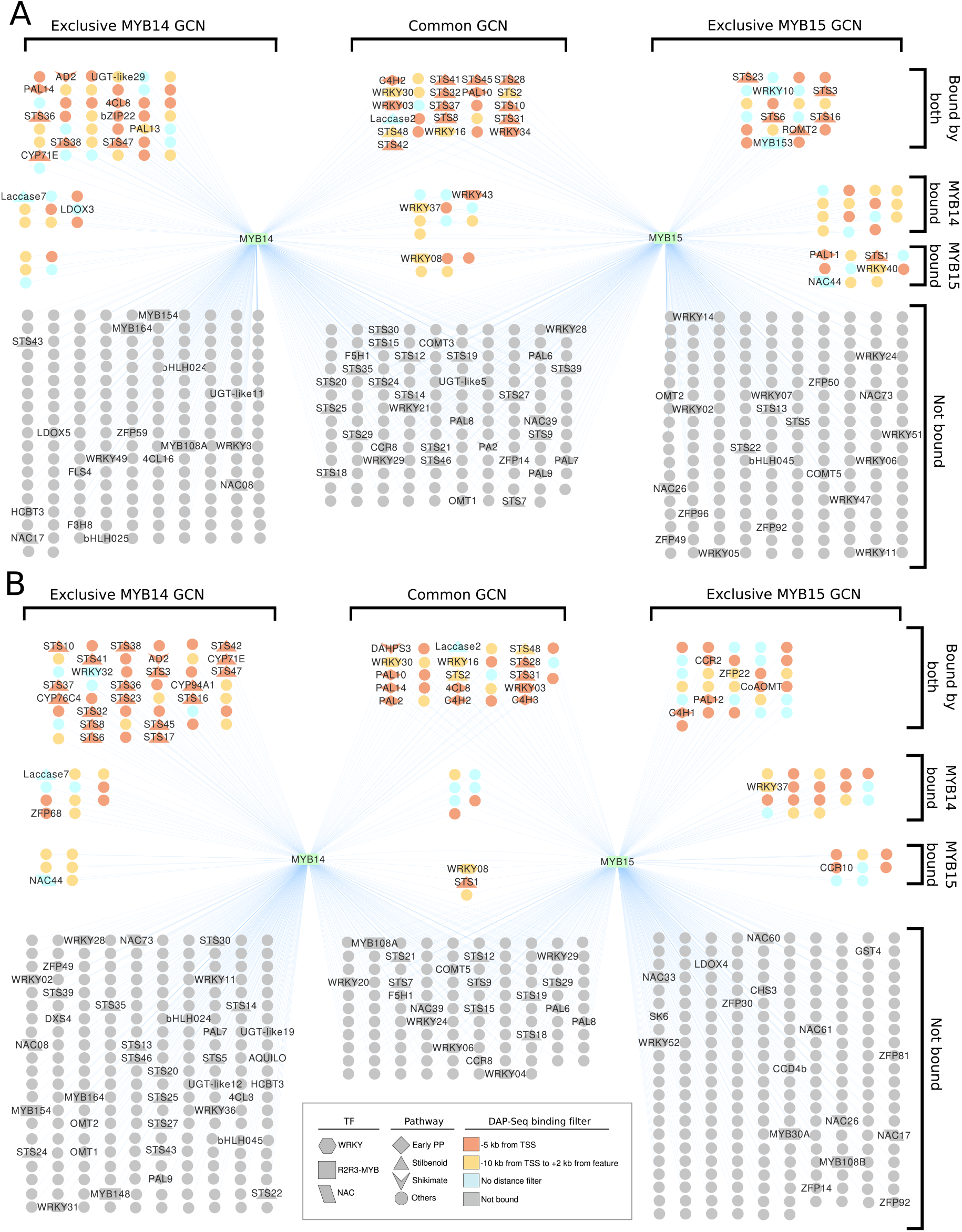
*MYB14* and *MYB15* leaf and fruit GCNs share many co-expressed genes, many of which are also detected with DAP-Seq. *MYB14* and *MYB15* GCNs consist of their top 420 co-expressed genes. The colour of each node (i.e. gene) depicts the different distance filters met by the closest DAP-Seq peaks. When genes are bound by both MYB14 and MYB15 the distance filters are only indicated if both TFs have peaks within them. The shape of each node determines the metabolic pathway or transcription factor family it belongs to. (a) Leaf GCNs. (b) Fruit GCNs.

### Integrating GCNs and DAP-Seq data to identify high confidence targets

In addition to the *MYB14* and *MYB15* GCNs, gene-centred GCNs were extracted for each DAP-Seq identified gene from the whole genome co-expression network (4,041 and 3,626 for *MYB14* and *MYB15* respectively). This allowed for further integration of co-expression and TF-binding results by overlapping DAP-Seq results with the newly extracted MYB-bound GCNs, as well as with each *MYB14* /*MYB15* GCN. For this overlap, only MYB-bound genes with peaks within 10kb upstream of the TSS and 2kb downstream of the end of their gene feature were considered.

Bound genes with a co-expression relationship present in at least one of the two GCNs (i.e. *MYB14* being present in a MYB14-bound gene GCN and/or *vice versa*) were considered as high confidence targets (HCTs). HCTs of both *MYB14* and *MYB15* were overlapped to obtain common HCTs. This integration of gene-centred GCNs increased the number of MYB-bound genes supported by network data. For instance, in fruit HCTs an increase from 76 to 145 and from 56 to 127 was observed for *MYB14* and *MYB15*, respectively. The different tissues used for the aggregate networks did not impact the total number of MYB HCTs, obtaining 146 and 145 *MYB14* HCTs in leaf and fruit, respectively. Gene enrichment analysis conducted for each MYB HCT list, using either Gene Ontology (GO) or Kyoto Encyclopedia of Genes and Genomes (KEGG), show trihydroxystilbene synthase activity as the most significant term (Fig. 4a and Dataset S2). Moreover, there are a number of interesting terms such as shikimate 3-dehydrogenase activity, phenylalanine ammonia-lyase activity and other shikimate/early phenylpropanoid related terms. Out of the 57 common fruit HCTs, many in fact correspond to *PAL* and *STS* genes. Although gene set enrichment analyses offer similar results for both tissue-specific HCTs, the defense response term was exclusive to leaf.

**Figure 4:**
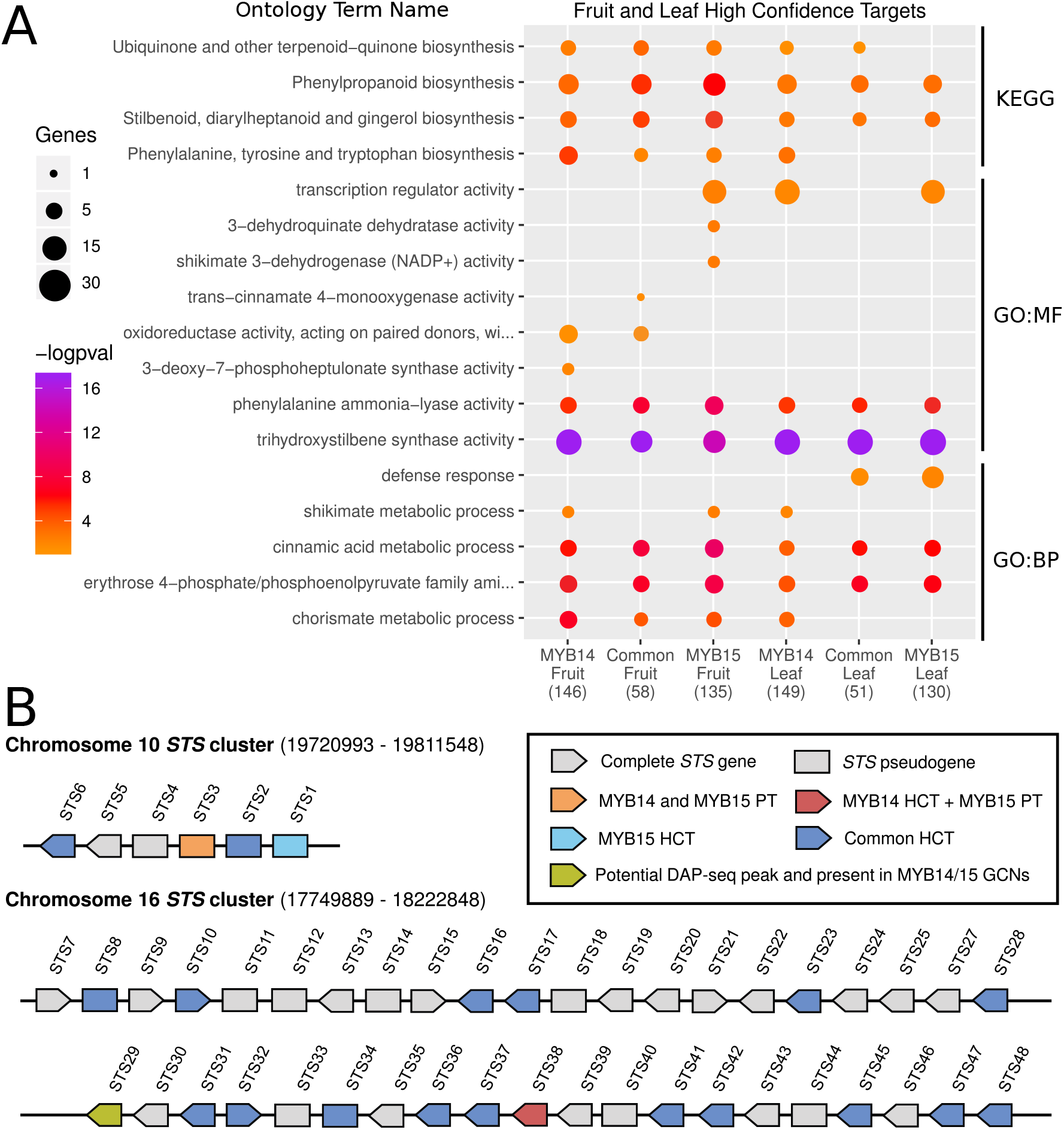
Gene set enrichment analysis of high confidence targets (HCTs) and *STS* gene synteny. (a) A selection of significantly enriched ontology terms related to secondary metabolism are shown for fruit and leaf HCTs (full lists in Dataset S2). A -logpval scale is provided where a higher value represents a greater statistical significance on a continuous colour scale from orange to purple. The number of targets intersecting with each ontology term is represented by point size. GO ontology levels are indicated as GO:MF (molecular function) and GO:BP (biological process). (b) *STS* genes cluster together in two different chromosomal regions, adapted from (Parage et al., 2012). DAP-Seq bound genes are referred to as putative targets (PTs) whilst HCTs (fruit or leaf) consider the integration of tissue-specific GCN and DAP-Seq data.

Common MYB14/MYB15 fruit HCTs show the presence of at least one known co-regulator of the stilbenoid pathway, i.e. *WRKY03*, as well as specific shikimate pathway enzymes, such as *DAHPS3* and *SDH4*, and key early phenylpropanoid enzymes such as *4CL8* and *C4H1-3*. Common fruit HCTs also contain 12 *STS* genes whilst individual fruit HCTs contain seven exclusive *STS* genes in the case of *MYB14* and one in the case of *MYB15*. Common MYB14/15 leaf HCTs show similar composition as fruit HCTs, with the presence of *WRKY03*, *4CL8*, *C4H2* and 17 different *STS* genes. Eight *STS* genes appear as HCTs for fruit and leaf considering both MYB14 and MYB15. Only one *STS* bound gene, the pseudogene *STS3*, is not a HCT (Fig. 4b). Compared to the many target genes within the shikimate, early phenylpropanoid and stilbenoid pathways a lack of HCTs is observed amongst the lignin or flavonoid branches of the phenylpropanoid pathway in the both fruit and leaf HCTs (Fig. S1, Dataset S2). To further corroborate some of the suggested MYB targets, we overexpressed *MYB15* in grapevine leaves and analysed their deferentially expressed transcriptomes.

### Transient *MYB15* overexpression in grapevine confirms many fruit and leaf predicted targets

We validated MYB15 high confidence targets by transiently overexpressing *MYB15* in grapevine leaves from 10 week old plants. Microarray analysis was conducted at 24, 48 and 96 hours after *in planta* agroinfiltration with a 35S:MYB15 construct or an empty vector control. Endogenous and total *MYB15* expression was monitored with respect to control samples showing a short-term increase at the initial time points probably due to the agro-infiltration *per se* (i.e. wounding stress). Nonetheless, a greater and longer-lasting increase in expression was observed thereafter in the *MYB15* -agroinfiltrated samples which can be fully attributed to the overexpression of the transgene (Figs 5a, S2). Gene expression data were normalised and clustered using weighted gene co-expression network analysis (WGCNA), resulting in 37 modules (Fig. 5b, left panel). We found the *MYB15* probe in module eigen-gene 24 (ME24), which is nearly identical to ME5, both showing a higher expression in 35S:MYB15 leaves compared to controls at all time-points. These two modules also clustered closely to ME27 which still showed similar z-score patterns across the different samples. By inspecting the number of genes present in leaf- and fruit-HCTs or MYB15-bound genes in each module and the clustering proximity to ME24, we determined that these four modules (ME24, ME5, ME27 and ME3) hold most of *MYB15* targets (Fig. 5b). The average gene expression of these four modules is indeed higher in *MYB15* -overexpressed samples with respect to control time-points (Fig. 5c). GO and KEGG enrichment analyses were carried out for each module (Fig. S3) revealing interesting enrichment terms such as the expected trihydroxystilbene synthase term. In addition, other terms were O-methyltransferase activity (ME5) where *ROMT1* gene is found, other shikimate/early phenylpropanoid enzymatic terms, and defense response terms such as “response to stress”, “response to fungus” and “response to chitin”. Probe-associated genes with positive fold changes cluster together and they mostly belong to the four modules of interest. Moreover, the probe representing *ROMT1* and 5 additional *ROMT* -like genes shows a strong fold induction across all timepoints (Fig. 5d). A similar induction dynamic is observed for other phenylpropanoid related enzymes such as 4CL8 and TFs such as WRKY03, WRKY08 or WRKY34. Conversely, a few bound genes with negative fold changes cluster together and belong to more distant modules.

**Figure 5:**
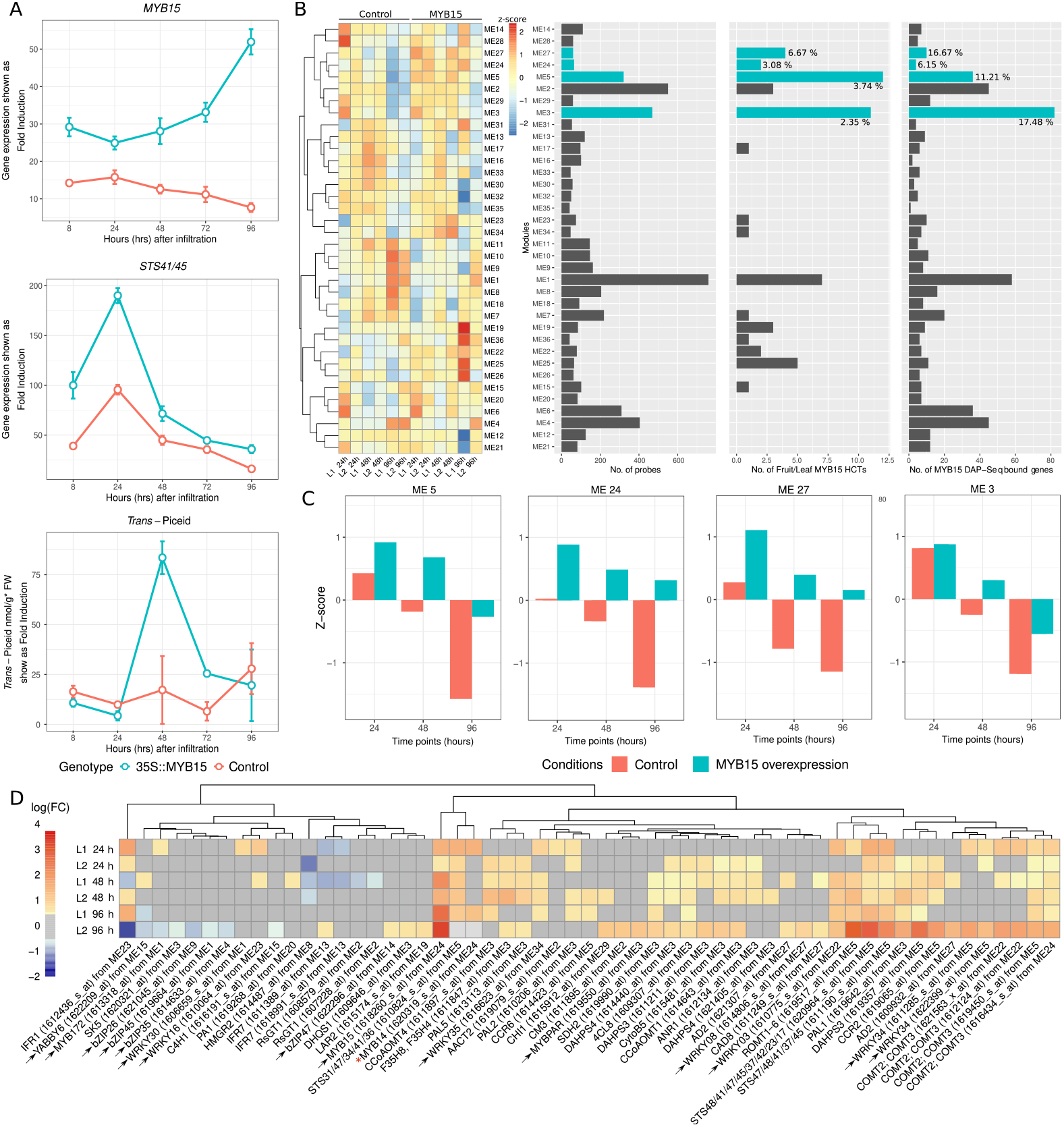
Secondary metabolism genes are induced upon overexpression of *MYB15* in grapevine leaves. (a) Increase in *MYB15* and *STS* expression as well as in *trans* -piceid content expressed as fold induction compared to non-infiltrated leaves for both 35S::VviMYB15 and empty vector control transformations (blue and red, respectively). (b) Clustered mean z-scores across each sample (leaf replicate and time point combination) for modules obtained by WGCNA from log(RMA+1) expression data. Barplots for every module show; i) the number of probes per module, ii) the number of probes with at least one MYB15 leaf or fruit HCT assigned and iii) the number of probes with at least one MYB15 DAP-Seq bound gene assigned. Percentages are provided for the four cluster modules of interest (marked in blue). (c) Mean z-scores across different leaf samples grouped by agro-infiltration treatment and time point for the four modules of interest. (d) Temporal gene expression changes in response to *MYB15* overexpression for MYB15-bound genes (as well as *MYB14* ) present in the grapevine reference gene catalogue (v1.1) with a Log_2_ fold change greater than 0.53 or smaller than -0.53 in at least one time-point. Log Fold change values between -0.53 and 0.53 are greyed out.

### *VviMYB13* shares a high proportion of bound genes with its two Subgroup 2 co-members

R2R3-MYB Subgroup 2 (S2) is composed of MYB13/14/15 in grapevine and Arabidopsis. Alignments and phylogenetic analyses of this subfamily in these and other plant species show that this close evolutionary relationship is in part explained by the conservation of the R2/R3 repeats and their C-terminal FW1 and FW2 domains (Fig. S4). The closest grape MYBs to S2 have lost at least one of these domains (i.e. *MYB135-138* ). Thus, we additionally checked the potential contribution of MYB13 to the regulation of secondary metabolism in grapevine. Our VviMYB13 DAP-Seq analysis reported 17,019 binding events assigned to 10,624 different genes (Fig. S5, Dataset S1). Most VviMYB14/15 DAP-Seq bound genes are contained within VviMYB13 bound genes (Fig. S5b). Around 74% of peaks are present between 10 kb upstream of transcription start sites (TSSs) and 2kb downstream of annotated gene ends. VviMYB13 binds to *STS* promoter regions in a similar manner to VviMYB14/15 (Fig. 6b, S6). The DNA binding motif obtained for VviMYB13 is practically identical to that of VviMYB14/15 (Fig. 1a) and even to the binding motif identified for AtMYB13 by O’Malley et al. (2016) (Fig. 6a). Interestingly, VviMYB13 was observed to bind at -2.3 kb from the TSSs of both *VviMYB13* itself and *VviMYB14*. Gene set enrichment analyses for AtMYB13 and VviMYB13/14/15 distance-filtered bound genes were carried out to illustrate the functional relatedness of these grape TFs in comparison to Arabidopsis (Fig. 6c and Dataset S1). Surprisingly, AtMYB13 also presents shikimate and early phenylpropanoid related terms as its *V. vinifera* homologues (e.g. phenylalanine ammonia-lyase activity). Since *A. thaliana* does not produce stilbenes, the trihydroxystilbene synthase activity terms is only significantly enriched for VviMYB13/14/15. An interesting term that is only present for AtMYB13 is lignin biosynthesis suggesting a potential regulaotry diversification between Arabidopsis and grape homologues.

**Figure 6:**
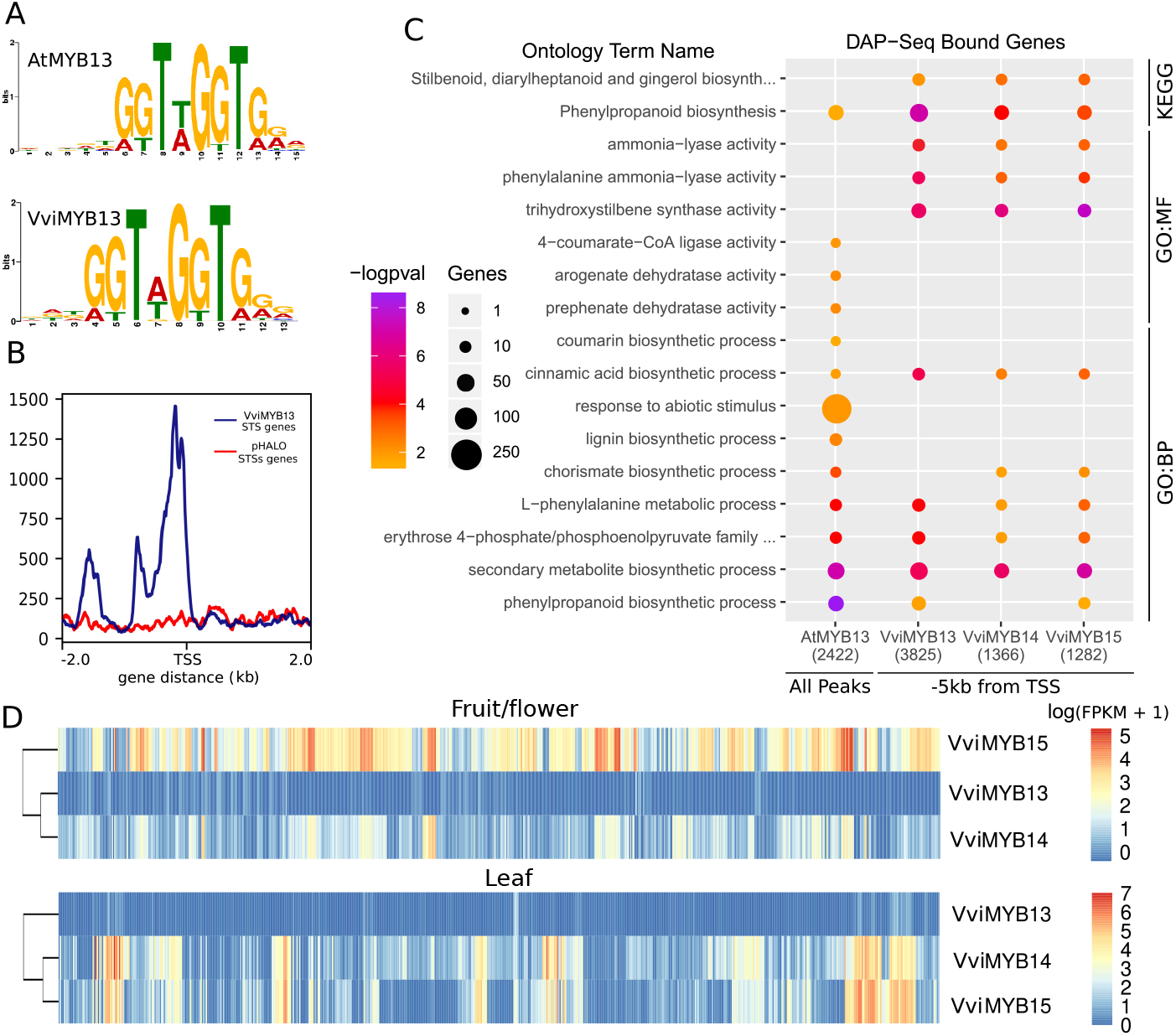
VviMYB13 binding features are similar to both VviMYB14 and VviMYB15 despite its different tissue-specific gene expression patterns. (a) AtMYB13 (O’Malley et al., 2016) and VviMYB13 DNA binding motifs identified using MEME suite. (b) VviMYB13 DAP-Seq binding peaks events with respect to 2 kb upstream and downstream of *STS* transcription start sites. The figure was generated using the Deeptools suite v.3.3.2, computing and normalising the coverage for each BAM file with a BinSize = 10 and RPKM normalisation. RPKM value for each bin is the average between the 10 positions that define each bin. BigWig files for the individual replicates for the TF were merged using bigWigMerge v.2 and bedGraphToBigWig v.4. (c) A selection of significantly enriched terms from KEGG and GO ontologies is shown for AtMYB13 and VviMYB13/14/15 bound genes. A -logpval scale is provided where a higher value represents a greater statistical significance on a continuous colour scale from orange to purple. The number of DAP-Seq bound genes intersecting with each ontology term is represented by point size. GO ontology levels are indicated as GO:MF (molecular function) and GO:BP (biological process). (d) A continuous colour scale from blue to red represents the log of FPKM+1 for *VviMYB13/14/15* across different SRA runs for every experiment used to build the fruit/flower and leaf aggregate networks.

Despite the similarities of VviMYB13/14/15 through DAP-Seq analysis, an overall lack of expression of *VviMYB13* across the SRA experiments used in the aggregate whole genome co-expression networks (thus, very low expression in leaves and flowers/fruits) meant that no valuable co-expression data could be extracted for *VviMYB13* (Fig. 6d). Transient overexpression of MYB13 in grapevine, or further inspection of condition-specific data (e.g. stress-related) where MYB13 is induced could be used to narrow down its bound genes to a list of HCTs.

As shown previously, DAP-Seq bound genes of VviMYB13/14/15 include many members of the shikimate pathway, the early phenylpropanoid pathway and the stilbenoid branch (Fig. S7). Surprisingly, we also found among bound genes those corresponding to the lignin and flavonoid branches, many of which are known to be oppositely expressed compared to *STS* and *MYB14/15* genes; as an example, chalcone synthase genes have been shown as strongly repressed under UV-C or fungal stress conditions which in turn largely activate *MYB14/15* and *STS* gene expression (e.g. (Vannozzi et al., 2012);(Blanco-Ulate et al., 2015)). This observation and the fact that these additional bound genes are not HCTs (Fig.S1) may suggest that they are negatively regulated by the interaction of other proteins with R2R3-MYBs from subgroup 2.

### Characterisation of new stilbenoid-pathway genes identified as Subgroup 2 MYB targets

Several MYB14-15 HCTs and MYB13-bound genes encode for different types of enzymes that have not been functionally validated yet. Among those potentially related to stilbene metabolism we found several o-methyltransferases, lacasses, hydroxylases (cytochrome P450s) and glycosyl-transferases, all representing interesting cases of further validation and which would position DAP-seq as a tool for novel enzyme identification.

As a first approach to study these potential novel enzymes of the stilbenoid pathway, we conducted a 7 day time-course experiment of grapevine (cv. ’Gamay Fŕeaux’) cell-cultures, elicited with methyl-jasmonate and cyclodextrins (MeJa+CD) that are known to largely and specifically activate the expression of *MYB/14/15* (Almagro et al., 2014). As cell suspensions from this ’*teinturier* ’ (i.e. red fleshed) cultivar ectopically accumulate anthocyanins in the presence of light, we grew and sub-cultured the cells for several passes in dark conditions to avoid the phenylpropanoid pathway to be prematurely committed for the production of flavonoids in detriment of stilbenoids being accumulated. Anthocyanin-devoid (i.e. white) cells were elicited for the quantification of secondary metabolites at four and seven days. An accumulation of resveratrol was observed only inside MeJA-treated cells at both time-points whilst anthocyanin content was greatly reduced compared to the control at the 7th day (Fig. 7; Fig. S8). Anthocyanins show an increasing tendency only in the control due to the effect of sugar replenishment (conducted at the beginning of the time-course) while in the elicited cells jasmonate seems to preferentially direct the flux of the phenylpropanoid pathway for stilbene accumulation rather than flavonoids despite of the effect of sugars. Moreover, there was an increase of piceatannol and viniferin (specially at day 7), while piceid accumulation did not present significant changes across treatment and time-points. We compared the accumulation of metabolites with the expression of the shikimate, early phenylpropanoid, and stilbene pathway genes (including the potential novel pathway genes) by reanalyzing the previously published microarray study of (Almagro et al., 2014) of 24h MeJA+CD-treated cv. ’Gamay Fŕeaux’ cells (Fig. 7. We first manually curated and improved the current MapMan grapevine ontology with the complete list of genes within the newly created stilbene pathway terms, amongst which we created “Trans-resveratrol di-O-methyltransferase activity” and “Resveratrol Glycosyltransferase activity” (Dataset S3). The combined DAP-Seq results of MYB13/14/15 were included in the context of these pathways, highlighting those gene that are bound by S2 MYBs. The metabolite profile observed in our experiment correlates with the activation of MYB15 and almost all its bound genes (*no probe for MYB14/MYB13* ), including resveratrol-modifying candidate genes. Production of viniferin and piceatannol also matched the up-regulation of their putative related enzymes. In particular, *Laccase8* and the cytochrome P450 *CYP76C4* genes, identified as MYB13/14/15 HCTs, could be directly responsible for the enzymatic reactions producing these compounds in our elicited grape cells. On the other side, the microarray data also supports the lack of pterostilbene accumulation, as no up-regulation of *ROMT1* or *ROMT-like* genes was observed.

**Figure 7:**
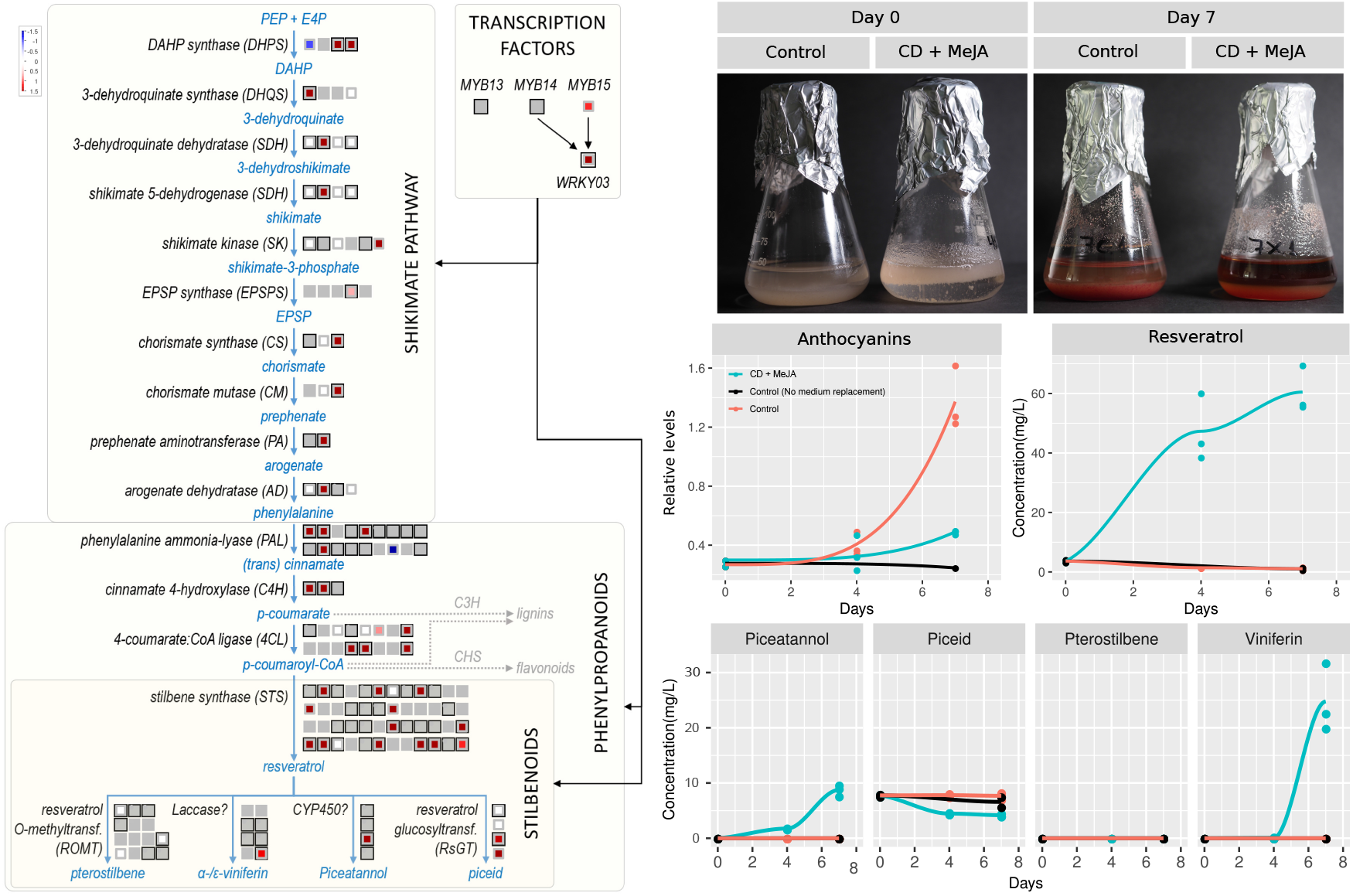
MYB13/14/15 DNA binding of shikimate, early phenylpropanoid and stilbenoid pathway genes shown together with microarray gene expression of methyljasmonate and cyclodextrins (MeJa+CD) elicited cells (Almagro et al., 2014). Left Panel: Genes are ordered from left to right and top to bottom. Genes that are surrounded by a black box correspond to MYB-bound genes (within -10kb of the TSS, the gene body itself, or +2kb from the end of the gene). Significant (0.05 p-value threshold) positive log2 fold-changes are shown in red and negative changes in blue. Genes with no associated microarray probe are greyed out whilst white boxes mark genes with no significant differential gene expression. *WRKY03* is included as it has been shown to cooperate with R2R3-MYBs for inducing *STS* gene expression ((Vannozzi et al., 2018)). Right Panel, Top: Grape cell suspensions at the beginning of the elicitation experiment (Day 0) and seven days after (Day 7). Bottom: Anthocyanin content, measured by spectophotometry, and stilbenoid quantification by LC-MS (i.e. resveratrol, piceatannol, pterostelbene, piceid and viniferin in the grape cells at Day 0, 4 and 7. An additional sample, corresponding to non-elicited grape cells sub-cultured in the old same growth media (i.e. with no sugar-replenishment) was taken at Day 7.

We inspected additional publicly available RNA-seq datasets to see if *MYB15/14* targets with potential resveratrol-modifying activity were co-expressed with their corresponding stilbenoid metabolites. For instance, by reanalyzing the transcriptome responses upon Botrytis infection stage 2 in cv. ’Semillon’ ((Blanco-Ulate et al., 2015)) we see a large group of potential resveratrol glycosyl-transferase (RsGTs), CYP450 and laccasses, many of which were identified as HCT, being highly induced (Fig. S9). In this case the expression of lacasses and RsGTs matched the differential accumulation of viniferins and piceid in response to infection.

As *ROMTs* transcripts or pterostilbene were undetected in the elicited-grape cells or in the Botrytis-infected samples, we further inspected their expression in other transcriptomics datasets to look for specific organs and developmental stages showing a condition-specific induction. The cv. ’Corvina’ atlas (Fasoli et al., 2012) represents a suitable dataset with more than fifty samples corresponding to all types of vegetative and reproductive tissues and green-to-mature stages. The stilbenoid pathway and the regulators *WRKY03* and *MYB14/15* show high expression in leaf and berry development, in particular at senescent or mature stages for both types of organs. For the case of *ROMT1* and *ROMT-like* genes we see an almost specific up-regulation in senescent leaves (Fig. S10). Thus, we extracted RNA from these senescent grapevine organs and amplified two *ROMT-like* genes, here named *ROMT2* and *ROMT3*. Together with the previously characterised *ROMT1* gene used as a positive control, we transiently overexpressed them in *Nicotiana benthamiana* leaves in combination with *STS48* that is also a MYB14/MYB15 HCT. All these three *ROMT* genes, which show several binding events upstream of their TSSs, were able to promote pterostilbene accumulation in tobacco leaves as seen by LC-MS analysis (Fig. 8). On the contrary, the sole overexpression of *STS48* only produced piceid accumulation while the agroinfiltrated empty vector didn’t produce any stilbenes. Taken altogether, the DAP-seq data presented here allowed us to select candidate pathways genes for their enzymatic characterisation *in planta*.

**Figure 8:**
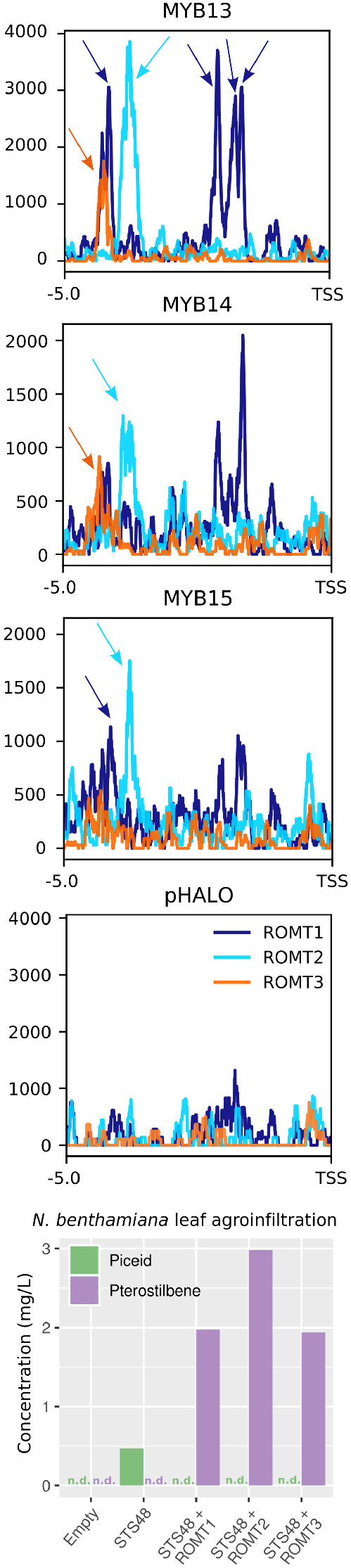
MYB13/14/15 DNA binding events in close proximity to *ROMT* genes. Peaks are shown in comparison to pHALO up to 5kb upstream of *ROMT1* -*3* TSSs. Detected peaks, marked by arrows, are found at around 4.5kb and 2kb upstream of TSSs. Agroinfiltration of both *STS48* and *ROMT1* -*3* leads to the accumulation of pterostilbene with respect to the empty vector control. Chromatograms and MS-detected transitions for piceid and pterostilbene are found in Fig. S11. Non determined metabolites due to very low concentration as marked as (n.d.).

## Discussion

### Gene co-expression and DNA-binding as a proxy for gene regulatory networks

Gene co-expression networks are based on the ”guilt-by-association” principle whereby correlation in expression implies biological association (Wolfe et al., 2005). This results in a promising tool for predicting gene regulatory networks, in particular to identify gene targets and their regulators. The constructed condition-dependent (i.e. leaf and fruit/flower) aggregate co-expression networks presented here take into account the frequency of particular gene-*to*-gene co-expression connections across independent transcriptomic experiments. On the other hand, non-aggregate networks just take into account the strength of particular gene-*to*-gene relationships across the complete set of biologically related samples from all experiments considered. Network aggregation has been shown to improve performance with the use of microarray grapevine datasets (Wong, 2020), and here it has been applied to all the leaf and fruit publicly available RNA-seq data for grapevine. The resulting network performance, estimated through the AUROC measurement (using MapMan BIN categories) was in our case `≈`0.75, which is comparable to previously published aggregate networks (Wong, 2020). Incorporating additional transcriptomic datasets is known to improve AUROC values until a plateau of approximately 0.8 is reached. This suggests that the number of experiments included in our final networks was enough to approach this value and hence support the derived co-expression connections amongst leaf- and fruit/flower-expressing genes. Nonetheless, a co-expression relationship does not indicate a biological connection in itself and the overlap with other datasets of experimental origin is required. This study represents a novel and successful application of gene co-expression networks in combination with genome-wide DNA-binding site interrogation used to draw the gene regulatory networks of transcription factors controlling specialised metabolism.

The three MYBs studied here belong to Subgroup 2 (S2) of the R2R3-MYB subfamily. Some members of this clade have been functionally characterised in other species. For instance, in *A. thaliana AtMYB13* is involved in response to UV-B (Qian et al., 2020), *AtMYB14* is related to cold tolerance (Chen et al., 2013) and *AtMYB15* is involved in the lignin biosynthesis and immune responses (Kim et al., 2020). In this study we show that the grape members of this subgroup are all tightly connected to the shikimate and phenylpropanoid pathways, specifically the early steps of the latter and most of the stilbenoid branch within this pathway. We show that not only *STS* genes are targets for these TFs but also resveratrol-modifying enzymes such as glycosyltransferases and o-methyltransferases, characterised for the first time here. We also suggest laccases and CYP450 hydroxylases as potential viniferin and piceatannol-producing enzymes based on DAP-Seq, co-expression networks and the inspection of large transcriptomic datasets where these genes are expressed in correlation with stilbenoid accumulation.

In this work we show a considerable overlap between MYB-related GCNs and their cistrome data, validating the use of co-expression relationships as a predictor of regulation. Non-overlapping genes present in GCNs may for instance represent indirect targets or genes commonly-regulated. In addition, many of the targets identified by the overlap of these datasets was also confirmed by the transcriptomic analysis of transient over-expressing plants. However, in the case of *MYB13*, due to its low expression in both leaf and fruit SRA experiments, we could not generate it as it would not have been very informative. Beyond target identification, GCNs can also point to MYB-interacting partners such as WRKY03 or WRKY43, which interact with MYB14 to increase *STS* expression (Vannozzi et al., 2018). Our GCNs also show a high co-expression of S2 MYBs with *MYB154* from S14 which has been recently shown to activate a few *STS* gene promoters in *Vitis quinquangularis* (Jiang et al., 2021).

### Shared and exclusive regulatory features of S2 MYB transcription factors

MYB13, MYB14 and MYB15 share a high proportion of DAP-Seq bound genes as well as identical DNA binding motifs suggesting a partial overlap of their regulatory roles. This conserved motif has already been described as a MYB *cis*-regulatory element; in particular the AC-element motif (CACC[T/A]ACC) previously identified for *AtMYB15* in *Arabidopsis thaliana* (Romero et al., 1998). As described in (Kelemen et al., 2015) by using the yeast one-hybrid approach, Arabidopsis S2 R2R3-MYBs show a high degree of specificity towards the AC-element motif, as well as other subgroups such as S1, S3, S13 and S24. Within the stilbenoid pathway, we found that MYB-binding patterns, i.e. in terms of their sequence and positions, across individual *STS* genes were remarkably similar for these three R2R3-MYBs, providing strong evidence that the recent tandem duplications observed for the *STS* gene family involved not only the gene body but also, up to a certain extent, their upstream regions. MYB13/14/15 TFs bind in two distinct *STS* promoter elements; the first at 400 bp from the transcriptional start site (TSS) that is present in most *STSs*, and a second at 2000 bp which is only present for a subgroup of them. This observation suggests that *STSs*, despite their high similarity in sequence and gene structure, may possess gene-specific differences in their regulation. The presence of two binding sites in some of them also offers the possibility of synergistic or dual-regulation. MYB TFs regulating other specialised metabolic pathways have been known to interact with other proteins from different TF families such as bHLHs and WDR as well as other MYBs. According to Höll et al. (2013) MYB15 and MYB14 lack a bHLH-interacting motif within their R3 repeat which is present in other bHLH-interacting MYBs (e.g. those from S5 and S6) but it is unclear if they have dimerisation domains. Further studies could address if MYB14 and MYB15 homo- or heterodimerise. If this is the case the idea of simultaneous binding at different regions of the promoter gains strength.

Our gene co-expression networks may help to point out *in vivo* regulatory differences between TFs with otherwise identical *in vitro* DNA-binding profiles. This appears to be the case for MYB14/15 as the exclusive presence of 28 *STS* genes in the *MYB14* fruit network points to a very interesting MYB14-specific relationship for a subset of *STS* genes. This observed contrast in MYB14 and MYB15 co-expressed genes could be explained by promoter differences in MYB14/15 leading to separate regulation by upstream TFs. On the other hand, structural changes between the TFs, leading to distinct interactions with other co-regulators, could also explain this observation. Most *STSs* are also MYB14 and MYB15 fruit HCTs. The two previously reported *STS* targets of MYB14/MYB15 (Höll et al., 2013) were either HCTs (*STS41* ) or present in the *MYB14* /*15* GCNs (*STS29* ). *STS29* was not defined as a fruit or leaf HCT since no binding events were automatically called by the peak detection software. However, a potential peak is observed for the three MYBs when examining the mapped sequencing reads in the promoter region of the gene.

The enrichment analyses of leaf and fruit high confidence targets, resulting from the overlap of GCNs and DAP-seq data, corroborates the newly found connections of *MYB13* /*14* /*15* with shikimate and early phenylpropanoid genes. Several *PAL* genes appear as HCTs in addition to *STSs*, supporting the idea that MYB14 and MYB15 may have the ability to increase and favour the activity of the phenylpropanoid pathway for the formation of stilbenes, in detriment of lignins or flavonoids. In fact, we may suggest that MYB14/15 might be involved in the repression, direct or not, of chalcone synthases, which are direct competitors of *STSs* activity. We also suggest that these MYB TFs may rewire the whole cell metabolic grid and favour the increase of carbon flux into the phenylpropanoid pathway (PPP) by activating the shikimate pathway. The direct DNA-binding observed for at least one isoenzyme of every shikimate and early PPP step highlights the central character of the studied R2R3-MYB regulators in activating stilbenoid metabolism. Despite not being able to perform leaf or fruit GCNs for *MYB13*, we suggest that these three proteins are highly redundant in their regulatory potential. In line with the partial redundancy of MYB14 and MYB15, the GCNs also suggest regulation differences between these two TFs.

Transcription factors act in a hierarchical manner, with major orchestrators (i.e. regulators of regulators) and pioneering factors (i.e. those binding to DNA before chromatin remodelling) at the top hierarchy. Based on our data, MYB14 and MYB15 seem to be high in this hierarchy, not only because they are able to control a large portion of the shikimate/early phenylpropanoid and stilbene pathways but also because around 20 and 22% of their targets are known transcription factors, respectively. In fact, at least nine *WRKY* transcription factor genes are present among the common *MYB14* /*15* HCTs. WRKY TFs have been suggested as MYB co-regulators in the control of *STS* expression [(Wang et al., 2020), (Xi et al., 2014), (Xu et al., 2019)] and also in many plant defence responses to pathogens. Particularly, *WRKY03* is bound by both TFs and present in *MYB14* and *MYB15* leaf and fruit GCNs. This observation adds one more layer of complexity to the hierarchical regulation of MYB14, which requires *WRKY03* to highly increase *STS* gene expression (Vannozzi et al., 2018).

The data presented here suggest that grapevine S2 members act mainly as positive regulators of transcription, however, based on the reduction of gene expression of some bound genes upon *MYB15* induction, S2 MYBs could also be involved in negative regulation possibly requiring other co-repressors. An interesting example is observed amongst circadian rhythm-related genes. Given the known circadian behaviour of *STS* gene expression (Carbonell-Bejerano et al., 2014), and the fact that circadian rhythm appears as an enriched termed amongst HCT and DAP-Seq lists (Datasets S1 and S3) we further explored the potential relationship of *MYB15* with the core molecular components of the circadian clock. *MYB15* leaf overexpression led to a strong down-regulation of *LHY* (Dataset S3), one of the two components of the main oscillator within the system. Moreover, different components of the clock such as *LHY* itself, *PPR7a*/*b*, *TOC-like*, *GI* and *ELF4* are bound by either of *MYB13* /*14* /*15* (Dataset S1), which further suggest a direct connection with the circadian rhythm. RVE8-type MYBs have been previously shown to have roles regarding the control of circadian rhythm in *Arabidopsis thaliana* (Shalit-Kaneh et al., 2018), however, the involvement of R2R3-MYBs has not been previously described.

### Integration of DAP-Seq and expression data as a tool for the identification of novel stilbenoid pathway enzymes

By overlapping the different datasets generated in this work, we were able to identify as MYB targets several still-uncharacterised genes that represent novel structural genes of the stilbenoid pathway, such as laccases (i.e. multicopper oxidases potentially involved in resveratrol dimerisation), resveratrol hydroxylases, resveratrol O-methyltransferases (ROMTs) and resveratrol glycosyl-transferases (RsGTs). Within O-methyltransferases, *ROMT1* has been the only gene described to date as involved in the synthesis of pterostilbene (Schmidlin et al., 2008), and here we show that *MYB13* and *MYB15* bind within its 5kb upstream region. Additionally, we show that the three S2 MYBs bind in a similar region with respect to *ROMT2* / and *ROMT10* promoters. Despite none of these ROMTs appear in *MYB14* or *MYB15* fruit GCNs, some of them do appear in leaf GCNs. This is in line with the lack of expression of *ROMT* genes in fruit or flower tissues (Fig. S10), highlighting the importance of tissue-specificity when dealing with co-expression networks. In fact, the transient overexpression of *MYB15* led to an up-regulation of all *ROMT1* -*6* enzymes recognised by the microarray probes. Clustering of microarray data (Fig. 5) also showed that the expression profiles of the *ROMT* enzymes was very similar to *MYB15* and hence the gene set enrichment analysis of cluster module ME5, adjacent to the *MYB15* -containing module ME24, showed enrichment of the O-methyltransferase activity ontology term.

We also identified cytochrome P450 and laccase (multicopper oxidases) among MYB-targets. Both of these enzyme classes are suggested to take part in the hydroxylation and oligomerisation of resveratrol. This has not been experimentally demonstrated except for the production of piceatannol from resveratrol by a transiently expressed human cytochrome P450 in *Nicotiana bethamiana* (Martínez-Márquez et al., 2016). We identify particular genes from these two families which are highly induced in stilbenoid-accumulating conditions such as Botrytis infection (e.g. at Stage 2 of the infection; Fig. S9) as well as being present in S2 MYB GCNs or HCT lists. Nevertheless, a further functional demonstration of their role as stilbene pathway genes is required in future studies.

Online tools for the inspection of DAP-Seq data would certainly help in the identification of novel TF-targets for their selection and further characterisation. As with RNA-seq results, DAP-seq data can be easily visualised in Genome browsers. One example is JBrowse (Buels et al., 2016), a genomics and transcriptomics data visualisation platform that allows to organize data in tracks, including the sequence of a reference genome assembly, annotation tracks and those containing alignment results (e.g. BAM, CRAM, or BigWig files). Using this tool, we developed a public grapevine DAP-Seq visualisation application named DAP-Browse, present in the Vitis Visualization Platform (Fig. S12; VitViz; available at https://tomsbiolab.com/vitviz).

## Materials and Methods

### DNA affinity purification sequencing

Genomic DNA (gDNA) was purified from young grapevine leaves of cv. ‘Pinot Noir’ (clone PN-94) according to (Chin et al., 2016). gDNA library construction and DAP-Seq were performed as described in (Bartlett et al., 2017), with slight modifications. Briefly, the gDNA samples (800 ng/library) were sonicated into 200 bp fragments on a focused ultrasonicator (Covaris). The gDNA library then underwent end-repair, A-tailing and adapter ligation for subsequent Illumina-compatible sequencing. Successful library construction was verified by gel electrophoresis of sonicated gDNA and qPCR of adapter ligated fragments. *MYB14*, *MYB15* and *MYB13* were amplified from cv. ‘Pinot Noir’ and cloned into the pIX-HALO expression plasmid (TAIR Vector:6530264275) with the HALO-tag at the N-terminal. Clones were verified by XhoI digestion. The *MYB* -expression vectors were used in a coupled transcription/translation system (Promega) to produce MYB Halo tagged proteins. Protein production was confirmed by western blot using an anti-HaloTag antibody (Promega). HaloTag-ligand conjugated magnetic beads (Promega) were used to pull down the HaloTag-fused TFs. Pulled-down TFs were exposed to DAP-Seq gDNA libraries for MYB-DNA binding. 400 ng of gDNA library was used per DAP-Seq reaction. The bound DNA was then eluted and sequencing libraries were generated by PCR amplification. Sequencing was carried out using Illumina NextSeq 500 set up to obtain 30 million 1x75 bp single-end reads. The original sequence data were submitted to the Gene Expression Omnibus (GEO) database of the NCBI under the accession GSE180450. An empty pIX-HALO expression vector was used as a negative control to account for non-specific DNA binding as well as copy number variation at specific genomic loci. Two biological replicates were used for all experiments including the control.

### TF peak calling and motif analysis

The latest published genome assembly for *Vitis vinifera* cv. ‘PN40024’ is the 12X.v2 which is associated to V1 and VCost.v3 annotations, containing 29,971 and 42,414 gene models, respectively [(Jaillon et al., 2007), (Canaguier et al., 2017)]. Although the VCost.v3 annotation has been manually curated for the *STS* family, it also contains non-protein coding genes (such as lncRNAs and miRNAs) most of which have only been automatically annotated. Since the main focus of this work was to study the regulation of protein-coding genes in secondary metabolism (stilbenoid pathway in particular), a new annotation file combining all V1 gene models with the updated *STS* VCost.v3 gene models was created. The merged gff3 annotation file is available at http://tomsbiolab.com/scriptsandfiles.

DAP-Seq reads were mapped to the ‘PN40024’ 12X.v2 reference genome using bowtie2 (Langmead and Salzberg, 2012) version 2.0-beta7 with default parameters and post-processing to remove reads that have MAPQ scores lower than 30. Peak detection was performed using GEM peak caller (Guo et al., 2012) version 3.4 with the 12X.v2 genome assembly using the following parameters: “*–q 1 –t 1 –k_min 6 –kmax 20 –k seqs 600 –k_neg_dinu_shuffle”*, limited to nuclear chromosomes. The biological replicates were analysed as multi-replicates with the GEM replicate mode. Peak summits called by GEM were associated with the closest gene model in the custom annotation file using the BioConductor package ChIPpeakAnno (Zhu et al., 2010) with default parameters (i.e. NearestLocation). *De novo* motif discovery was performed by retrieving 200 bp sequences centred at GEM-identified binding events for the 600 most enriched peaks and running the meme-chip tool (Machanick and Bailey, 2011) in MEME suite 4.10.1 with default parameters.

### Generation of condition-dependent aggregate whole genome co-expression networks and extraction of gene-centred co-expression networks

Transcriptomic RNA-Seq Sequence Read Archive (SRA) studies (Illumina sequencing) from fruit/flower and leaf tissue samples were downloaded. SRA studies were manually inspected and filtered to keep those which were correctly annotated and contained four or more data sets (i.e. runs) whilst also excluding SRA studies concerning microRNAs, sRNAs and non-coding RNAs. A total of 35 and 42 SRA studies were obtained, encompassing 807 and 670 runs from fruit/flower and leaf samples, respectively (Dataset S2).

Single- and paired-end runs were trimmed separately using fastp (Chen et al., 2018), version 0.20.0 with the following parameters: *“–detect adapter for pe –n base limit 5 cut front window size 1 cut front mean quality 30 –cut front cut tail window size 1 cut tail mean quality 30 –cut tail -l 20”*. After the trimming, the runs are aligned with STAR (Dobin et al., 2012), version 2.7.3a, using default parameters. Raw counts were computed using FeatureCounts (Liao et al., 2013), version 2.0.0, and the VCost.v3 27.gff3 (available at http://www.integrape.eu/index.php/resources/genomes/genome-accessions) gene models. Each SRA study was individually analysed to build a highest reciprocal rank (HRR) matrix (Mutwil et al., 2011). Briefly, raw counts for every run within the same SRA study were summarised in individual count matrices. Each raw counts matrix was normalised to FPKMs, and genes with less than 0.5 FPKMs in every run of the SRA study were removed. The Pearson’s correlation coefficients (PCC) of each gene against the remaining genes was then calculated for each SRA study and ranked in descending order. Ranked PCC values were used to compute HRRs amongst the top 420 ranked genes (420 roughly equals 1% of all VCost.v3 gene models), using the following formula: *HRR(A, B) = max(rank(A,B), rank(B,A))*, generating a HRR matrix for each SRA study. To construct the aggregate whole genome co-expression network, the frequency of co-expression interaction(s) across individual HRR matrices was used as edge weights, and after ranking in descending order, the top 420 frequency values for each gene were chosen to build the final aggregate networks.

Network functional connectivity (i.e. performance) across all given annotations and genes was assessed as in (Wong, 2020) by neighbour-voting, a machine learning algorithm based on the guilt-by-association principle, which states that genes sharing common functions are often coordinately regulated across multiple experiments (Verleyen et al., 2014). The evaluation was performed using the *EGAD* R package (Ballouz et al., 2016) with default settings. The network was scored by the area under the receiver operator characteristic curve (AUROC) across MapMan V4 BIN functional categories associated to the VCost.v3 annotation (Dataset S2) using threefold cross-validation. MapMan BIN ontology annotations were limited to groups containing 20–1000 genes to ensure robustness and stable performance when using the neighbour-voting algorithm. The top 420 most highly co-expressed genes for any gene of interest, was used to generate individual gene-centred co-expression networks (GCNs).

### Prediction of high confidence targets in grape reproductive and vegetative organs

To identify *MYB14* and *MYB15* high confidence targets (HCTs), TF-bound genes identified with DAP-Seq were overlapped with data extracted from the aggregate whole genome co-expression network derived from fruit/flower and leaf transcriptomic data. Individual GCNs were extracted from the whole genome network for each MYB14/15 bound gene and also for both *MYBs* to check if a particular *MYB* -*to*-target gene pair had at least one co-expression relationship. This relationship was considered positive if either the DAP-Seq bound gene was present in the *MYB* GCN or the respective *MYB* gene was present in the GCN of the candidate gene. Thus, DAP-Seq bound genes for each MYB TF with a positive co-expression relationship with their respective MYB TF were considered as fruit/flower or leaf HCTs. Gene Ontology (GO) and Kyoto Encyclopedia of Genes and Genomes (KEGG) enrichment analyses for HCTs were performed using the gprofiler2 R package (Kolberg et al., 2020) with default settings. A significance threshold of 0.05 was chosen for p-values adjusted with the Benjamini-Hochberg correction procedure (Benjamini and Hochberg, 1995).

### Transient *MYB15* overexpression, microarray time-course and *trans*-piceid quantification in leaves

*V. vinifera* cv. ‘Shiraz’ leaves from 10 week old plants were transformed via *Agrobacterium*-mediated infiltration as in (Merz et al., 2015) with *MYB15* (35S::VviMYB15) or an empty vector control. A GFP vector (35S::GFP-GUS) was used in each Agro-infiltration to estimate transformation efficiency. Samples were subsequently collected at five different time points (8h, 24h, 48h, 72h and 96h) in separate replicate pools (L1 and L2) consisting of three leaves from independent plants per time-point. Microarray analysis, using the Affymetrix *V. vinifera* Grape Array (A-AFFY-78), was carried out on extracted leaf RNA from three of the time points described for both control and *MYB15* infiltrated samples. Affymetrix probe sequences (originally associated to V1) were re-aligned using bowtie with default parameters against the ‘PN40024’ transcriptome derived from the VCost.v3 annotation to improve V1 probe-*to*-gene assignments (Dataset S3). Since each Affymetrix probe has a different number of sequences associated to it (ranging from 8 to 20), a probe-*to*-gene association was accepted when more than 40% of the probe’s sequences matched the same gene with no mismatches. The percentage was increased to 60% for hits with 1 mismatch, 70% with 2 mismatches and 80% with 3 mismatches. From a total of 16602 probes, 59.8% were assigned to a VCost.v3 gene model. The LIMMA (Ritchie et al., 2015) R package was used for RMA normalisation of fluorescence values after which they were log_2_-transformed. Probes with log_2_-fold-change values between -0.53 and 0.53 (corresponding to an absolute fold change of 1.45) for all *MYB15* overexpression time points were filtered out, obtaining 5,593 probes corresponding to deferentially expressed genes. RMA values of the filtered probes were clustered applying the weighted gene co-expression network analysis (WGCNA) R package (Langfelder and Horvath, 2008) using blockwiseModules with a soft-threshold power value of 18 (as no free-scale topology was reached), a deepSplit of 4 and a mergeCutHeight of 0.1, obtaining a total of 36 modules for the signed network (i.e. considering the sign of correlation). The pheatmap R package was used to represent the modules using kmeans clustering and the calculated z-scores across samples. The probe-assigned genes in each module were used for GO and KEGG enrichment analyses as described for HCTs. Quantitative expression analysis of *MYB15* or *STS* genes and *trans*-piceid quantification were conducted using L2 leaf pools across the different samples. *Trans*-piceid was quantified on a reverse phase HPLC instrument (Kronton). Quantitative PCR was run in triplicate per time-point and treatment using primers against the endogenous (aligning to the UTR) and transgenic *MYB15*, *STS25/27/29* and *STS41/45*. Transcript levels were corrected using the housekeeping gene *Ubiquitin1*. All used primers are shown in Dataset S3. The fold of induction reported for both mRNAs and metabolites was calculated relative to background levels in non-infiltrated leaves.

### Grapevine cell culture elicitation time-course

Liquid cell suspensions were initiated by transferring solid calli into a 250ml flasks containing liquid culture media (with Gamborg basal salts and Morel vitamins) described in (Bru et al., 2006) and grown in an orbital shaker at 120 rpm. Liquid cell cultures were maintained in dark conditions for two months, with subcultures being performed every 2 weeks by moving half of the extracellular volume to a new flask and restoring the original volume with fresh media (half medium replenishment). For the elicitation time-course, four flasks of cell suspensions were mixed together and 3/4 of the suspension was filtered using a sterile glass filter (90-150 µm pore size) and a vacuum filtration system to separate the cells from the media. A total of 15 100ml flasks were prepared with three biological replicates for each time-point and treatment. Day 0 cells were immediately collected after filtering and frozen in liquid nitrogen. For preparing Day 4 and Day 7 samples, 8 g of filtered cells and 32 g of new liquid media were added in each flask (complete medium replenishment). Treated (elicited) cells were grown in the liquid media with 50 mM methyl-beta-cyclodextrin (CAS No. 128446-36-6) and 84 µl of 47,5mM methyl-jasmonate dissolved in 100% MeOH while untreated (control) cell suspensions were grown in liquid media with 84 µl of 100% MeOH. The rest of the original suspension (1/4) was sub-cultured as usual and kept growing at dark for being used as an additional control for half medium replenishment at Day 7. All the replicates were grown and kept at 120 rpm in the conditions described above. Cell cultures were sampled at their corresponding time-point, filtered using a vacuum-aided system and washed with cold sterile water before freezing liquid nitrogen. Cells were lyophilised at -52°C and 0.63 mbar for 48 hours and then transferred to -80° C.

### STS and ROMT overexpression in tobacco leaves

*ROMT2-3* were amplified using TOPO-D (Invitrogen) compatible primers from cDNA derived from grapevine senescent leaves. RNA extraction and cDNA synthesis were performed using the Spectrum™ Plant Total RNA Kit (Sigma-Aldrich) and NZY First-Strand cDNA Synthesis Kit, respectively. The final *ROMT2-3* -containing pENTR/D-TOPO plasmids were verified by Sanger sequencing. *ROMT2* and *ROMT3* were transferred into the binary destination vector pB2GW7 including Cauliflower mosaic virus (CaMV) 35S promoter through Gateway LR clonase recombination (Invitrogen). Together with *ROMT1* and *STS48* constructs (Schmidlin et al., 2008; Santos-Rosa et al., 2008) all these were chemically transformed in *Agrobacterium tumefaciens* strain C58 for subsequent agroinfiltration of *Nicotiana benthamiana* leaves. The bacterial suspension was kept at an OD600 of 0.2 for each construct before infiltration of leaves on their abaxial side (three leaves of one plant for each experimental combination) using a 1 ml syringe without a needle. Agroinfiltration was conducted with the individual empty vector-, STS- or combined STS and ROMT-containing bacterial cultures. After 72h, leaf-tissue samples were collected by dissection so as to avoid leaf nerves. Collected samples were frozen at -80° C.

### Stilbenoid and anthocyanin quantification of grape cell cultures and Nicotiana agroinfiltrated leaves

Stilbenoid metabolites were extracted using 10 mg of cells in 1ml of 80% methanol with 220 rpm overnight shaking at 4°C. The extracted solution was centrifuged at 14.000 x g for 10 minutes, and collected in another tube for HPLC-MS detection and quantification. For the case of tobacco samples, leaf samples were weighed and grounded in liquid nitrogen. Extraction using 100% MeOH was carried out keeping a ratio of 8 ml of MeOH per 1 g of fresh tissue with 220 rpm overnight shaking at 4°C as described in (Martínez-Márquez et al., 2016). After 10 min centrifuging at 14.000 x g the supernatant was used for HPLC-MS determination and quantification. Anthocyanin quantification was performed as described in (Nakata and Ohme-Takagi, 2014). Briefly, lyophilised cells (o.2 ml) were treated with 1 ml of extraction buffer (45% MeOH and 5% HCL). After a 10 min vortex and centrifugation at 12.000 x g for 5 minutes, the supernatant was collected into a new tube and centrifuged again under the same conditions. Supernatant absorbance was measured at 530 nm and 637 nm and the anthocyanin relative levels were calculated using the formula: *(Abs*_530_ *–(0.25*Abs*_637_*))*5/(mg of cells)*.

### Re-annotation of the grapevine MapMan ontology and representation of previously published microarray and RNA-seq data

MapMan v4 (Schwacke et al., 2019) annotation was generated based on EGAD, resulting in high-quality annotations for 13,318 out of 41,413 VCost.v3 genes (32%) (Dataset S3). A pathway image for shikimate, phenylpropanoid and stilbenoid pathway was created and annotated using the MapMan tool (Thimm et al., 2004) for both Vcost.v3 and V1 of the grapevine genome annotations. This image includes the DAP-seq results whether a gene bound by MYB14, MYB15 and MYB13 is shown in black squares. All files are available for download from the GoMapMan exports page (Ram̌sak et al., 2014) (gomapman.nib.si/export). Significant Log_2_ fold-change values from a microarray study involving jasmonate elicitation of cv. ’Gamay Fŕeaux’ cell cultures (Almagro et al., 2014) and from a RNA-seq study of Botrytis infected berries of cv. ’Semillon’ ((Blanco-Ulate et al., 2015)) were represented within the newly generated MapMan pathway image.

### Funding and Acknowledgements

This work was supported by Grant PGC2018-099449-A-I00 and by the Ramón y Cajal program grant RYC-2017-23645, both awarded to J.T.M., and to the FPI scholarship PRE2019-088044 granted to L.O. from the Ministerio de Ciencia, Innovación y Universidades (MCIU, Spain), Agencia Estatal de Investigación (AEI, Spain), and Fondo Europeo de Desarrollo Regional (FEDER, European Union). C.Z. is supported by China Scholarship Council (CSC) no. 201906300087. K.G. and Z.R. were supported by the Slovenian Research Agency (grants P4-0165 and Z7-1888). S.C.H. is partially supported by National Science Foundation grant PGRP IOS-1916804. This article is based upon work from COST Action CA 17111 INTEGRAPE, supported by COST (European Cooperation in Science and Technology). Data has been treated and uploaded in public repositories according to the FAIR principles, in accordance to the guidelines found at INTEGRAPE. The genomic data presented here will be re-analysed and associated to the PN40024.v4 assembly and its structural annotation upon its release. Grapevine cell cultures in solid media were kindly provided by Roque Bru (Universidad de Alicante). Special thanks to Pere Mestre and Philippe Hugueney (INRAE Colmar) for providing the 35S:*STS48* construct and the 35S:*ROMT1* positive control used for agro-infiltration, and to Anne-Francoise Adam-Blondon and Nicolas Francillonne (URGI INRAE Versailles) for guidance in the adaptation of JBrowse during an INTEGRAPE Short-Term Scientific Mission of L.O.

## Supporting Information

**Figure S1:**
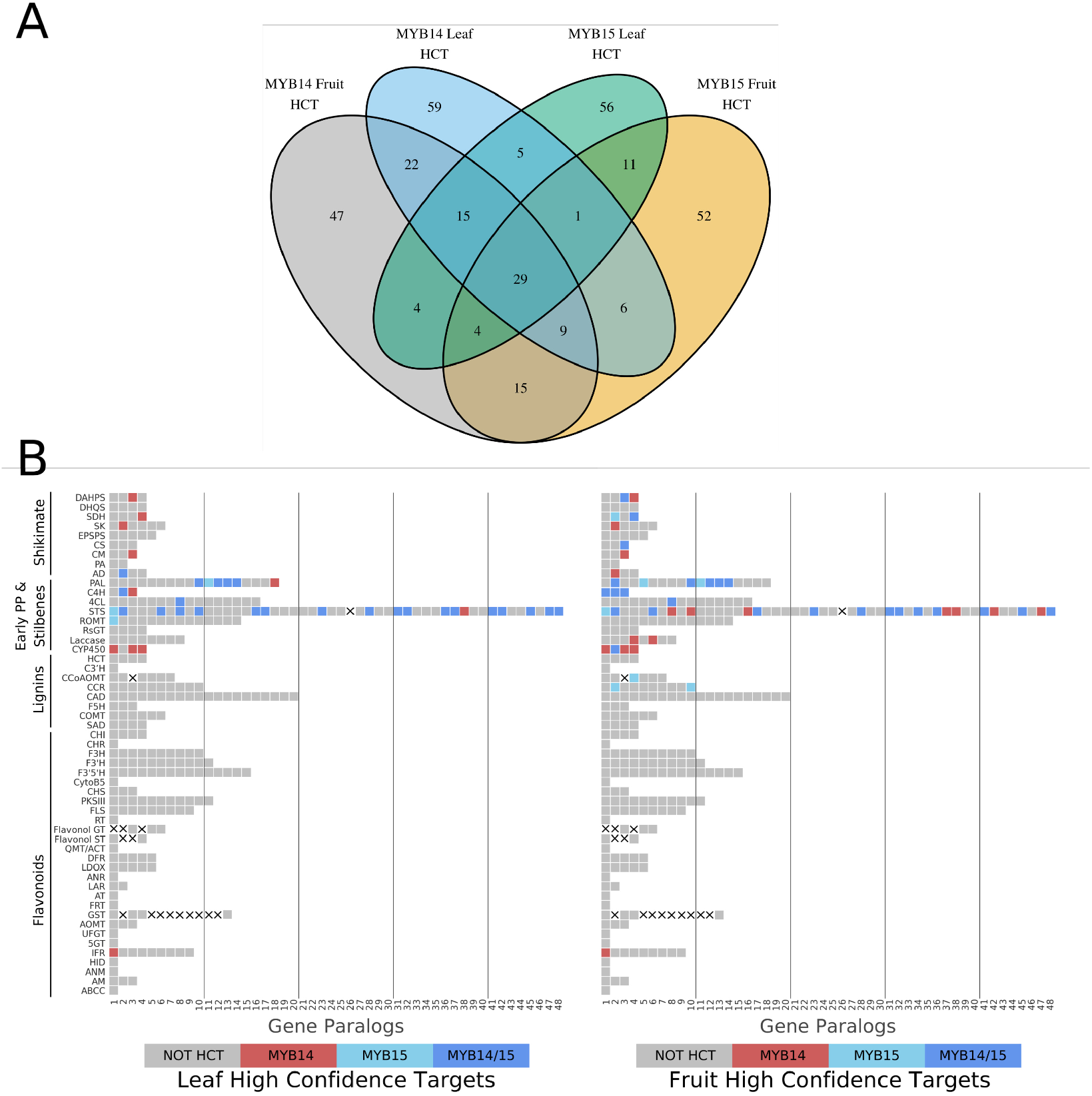
*MYB14* and *MYB15* High Confidence Targets (HCTs) amongst shikimate, phenylpropanoid and stilbenoid pathway genes. The x axis shows different gene numbers (e.g. isoenzymes). A cross marks gene symbols which do not exist (e.g. *STS26* is now part of *STS25* ) or are not associated to the respective pathways. *MYB14* HCTs are shown in red, *MYB15* HCTs in light blue and common HCTs in a darker blue.

**Figure S2:**
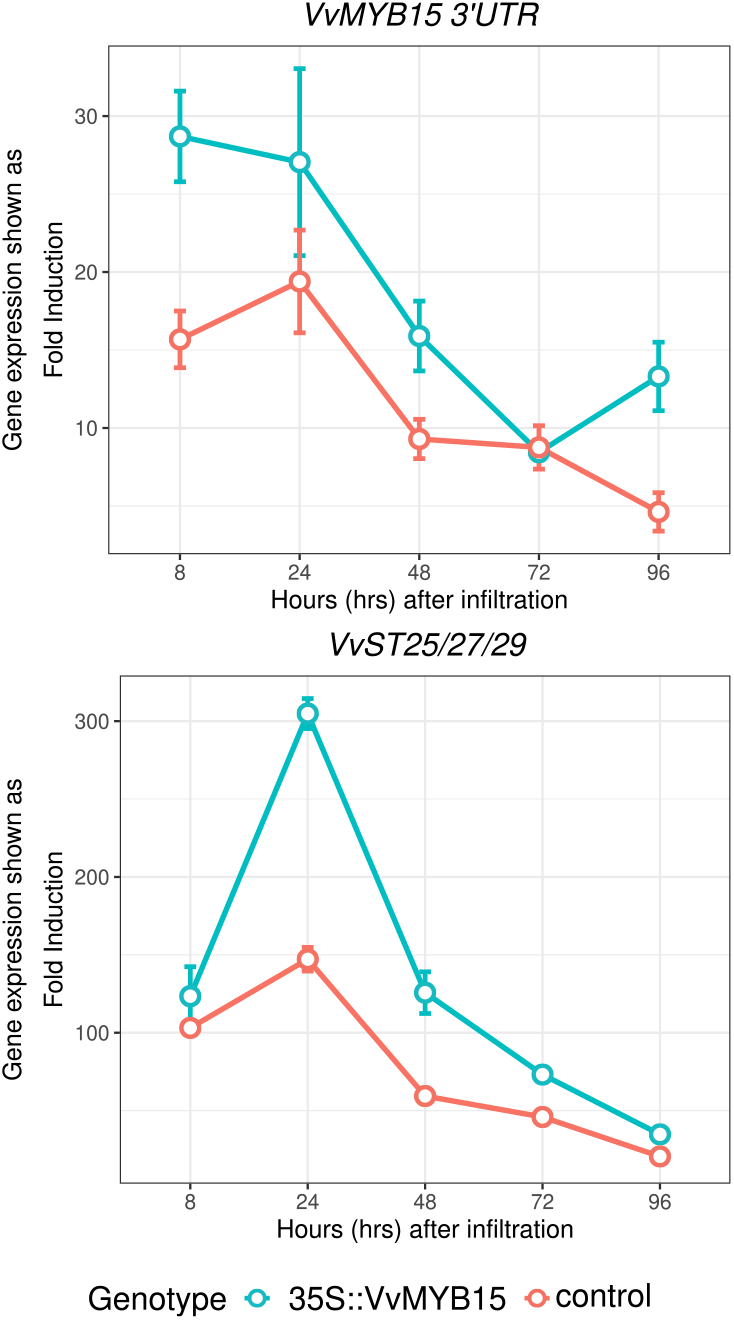
Endogenous *MYB15* and *STS25* /*27* /*29* expression in grapevine leaves overexpressing *MYB15*. Endogenous *MYB15*, shown by amplification with MYB15 3’UTR primers, is induced similarly with both 35S::VvMYB15 and empty vector control agro-infiltration. *STS25* /*27* /*29* expression is induced more with 35S::VvMYB15 agro-infiltration.

**Figure S3:**
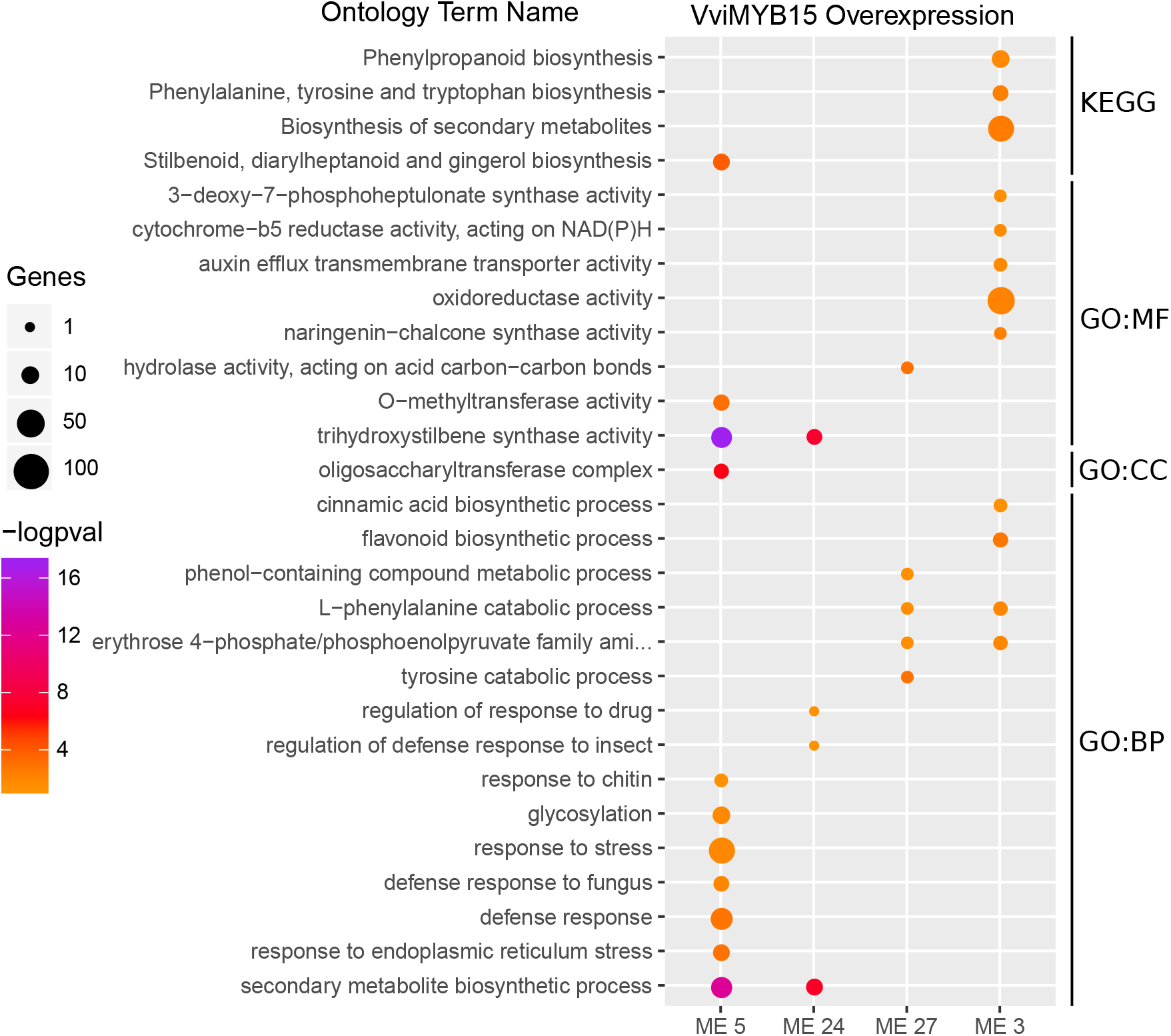
GO and KEGG gene set enrichment analysis for selected cluster modules in grapevine leaves overexpressing *MYB15*. A selection of relevant ontology terms are shown for the cluster modules of interest. A -logpval scale is provided where a higher value represents a greater statistical significance on a continuous colour scale from orange to purple. The number of genes intersecting with each ontology term is represented by point size. GO ontology levels are indicated as GO:MF (molecular function), GO:CC (cellular component) and GO:BP (biological process).

**Figure S4:**
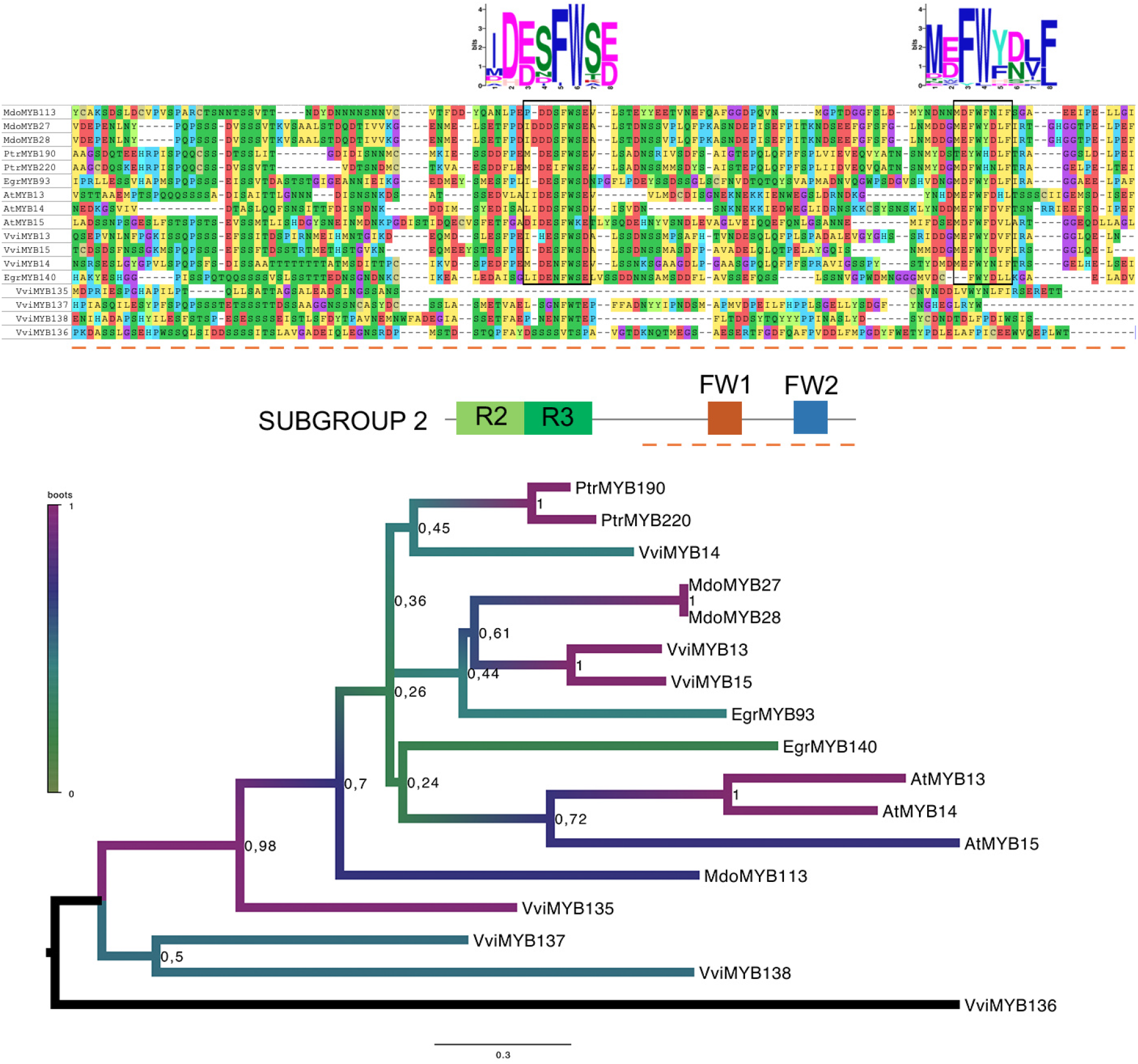
Phylogenetic and protein motif analyses of Subgroup 2 R2R3-MYB proteins and their closest relatives. (Top) Multiple protein alignments of the C-terminal domains of S2 members, including FW1 and FW2 conserved motifs. (Bottom) Rooted maximum likelihood (ML) phylogenetic tree constructed using complete protein sequences (1000 iterations). Evolutionary distances are represented as amino acid substitutions per site. Bootstrap values are shown on a coloured scale ranging from green to purple. Sequences from other species such as *Arabidopsis thaliana* (At), *Malus x domestica* (Mdo), *Populus trichocarpa* (Ptr) and *Eucalyptus grandis* (EGr) are shown for evolutionary context.

**Figure S5:**
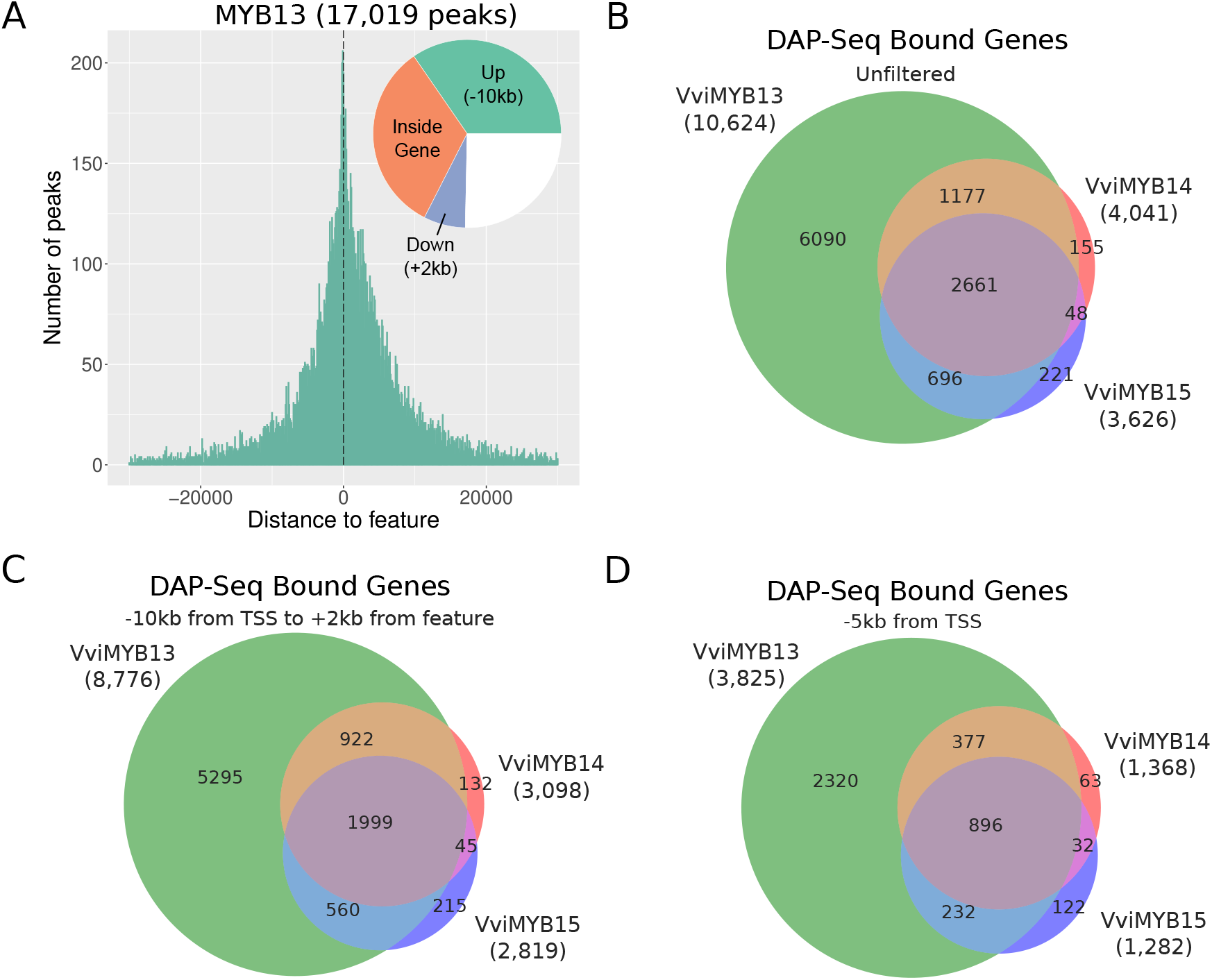
MYB13 DAP-Seq binding peak distribution and an overlap of MYB13/14/15 bound genes. (a) DNA binding events of MYB1 with respect to all transcription start sites (TSSs) of assigned genes. The proportion of binding peaks 10kb upstream of TSSs, inside genes or 2kb downstream of genes are represented within the pie-charts in green, orange and blue, respectively. (b-d) Overlap of MYB13, MYB14 and MYB15 DAP-Seq bound genes (in green, red and blue, respectively) with the following distance filters; no filter, a -10kb from TSS up to +2kb after end of feature and a -5kb from TSS filter.

**Figure S6:**
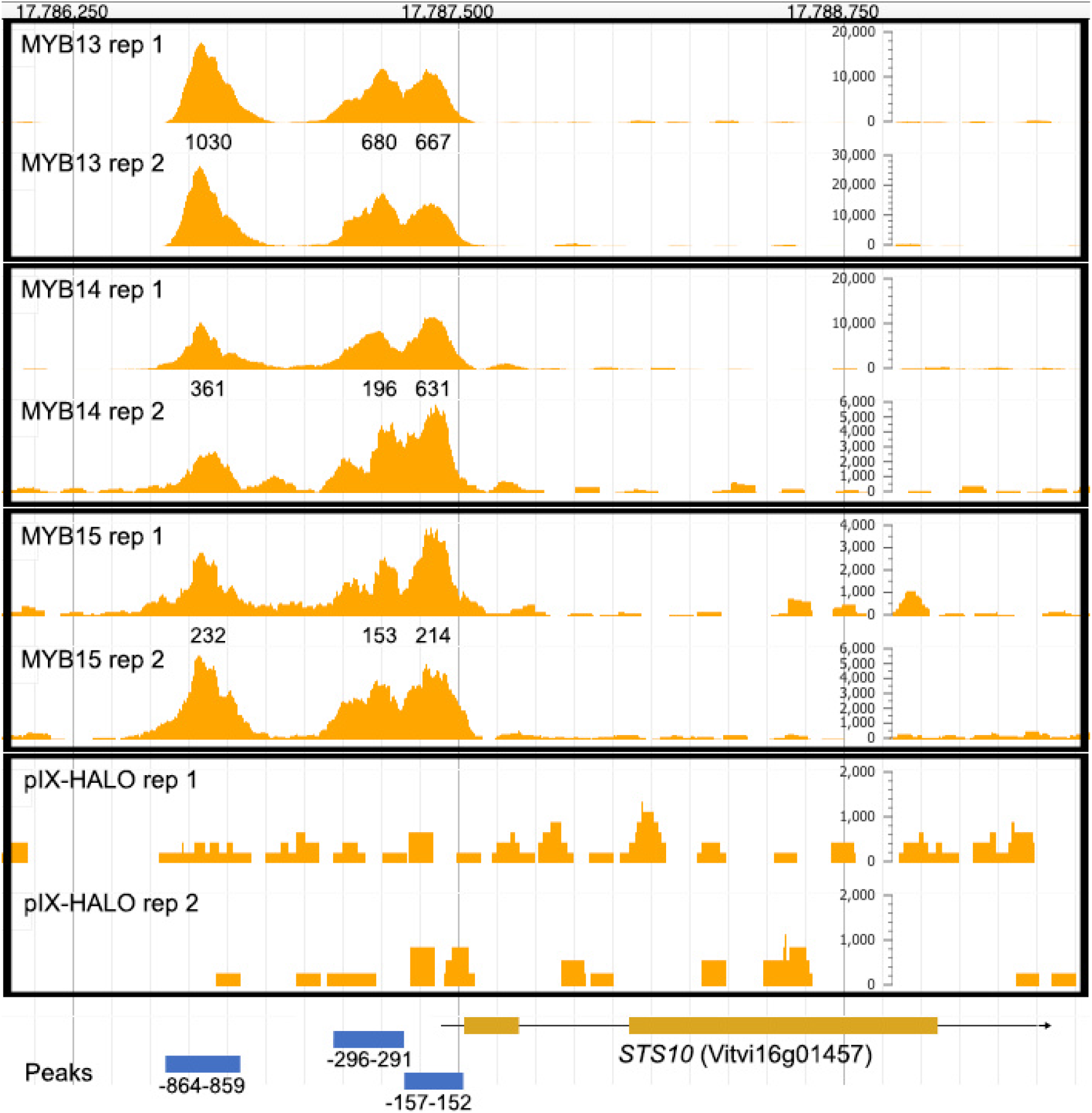
Genome Browser screenshot of MYB13/14/15 DAP-Seq detected peaks for *STS10*. JBrowse screenshots are assembled to show read alignments in the proximal promoter region of *STS10* for individual replicates. The three DAP-Seq detected peak regions are indicated in blue, whilst peak signal values obtained from combined replicates are indicated for each TF binding peak. The pIX-Halo control illustrates non-specific DNA binding.

**Figure S7:**
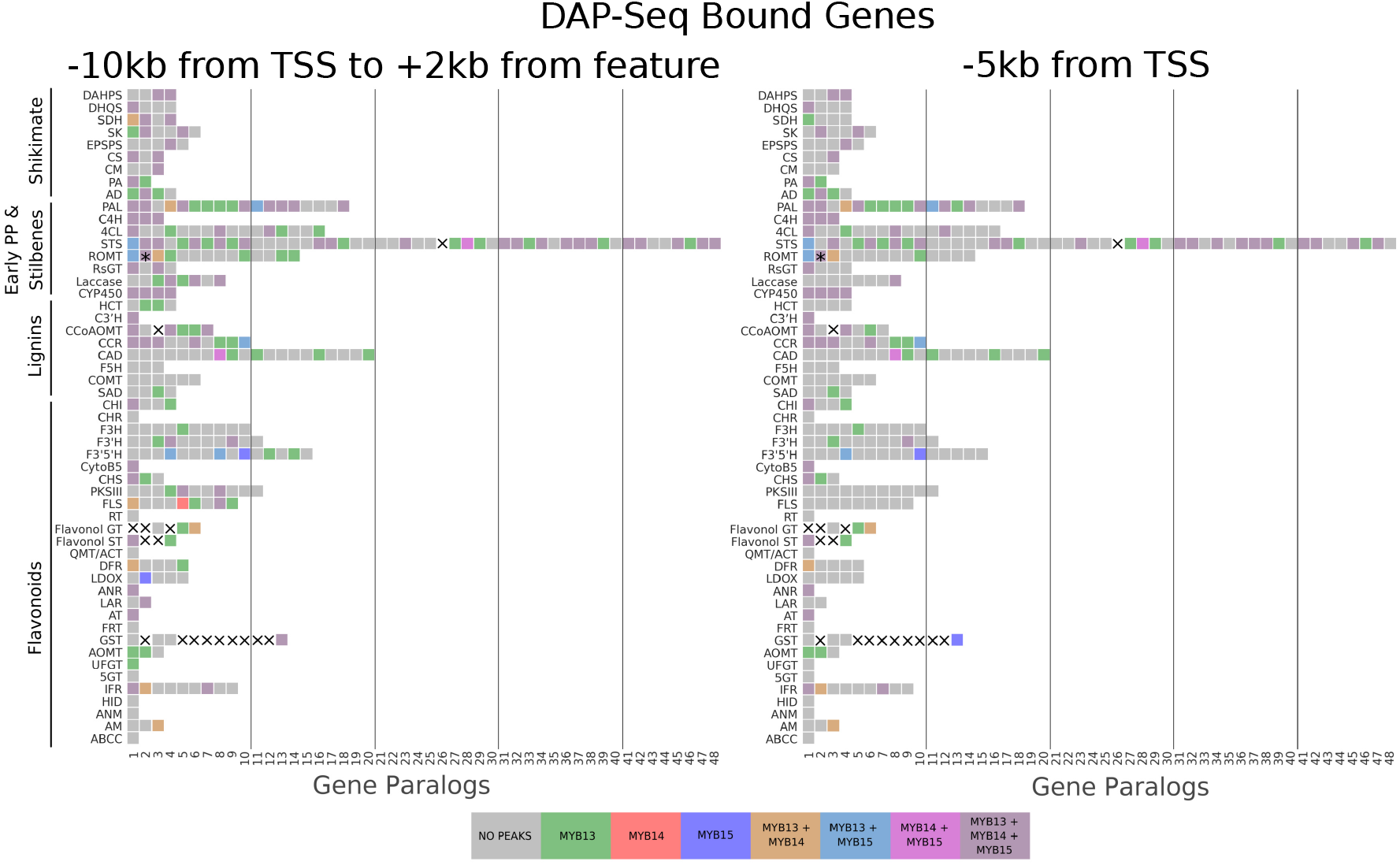
MYB13, MYB14 and MYB15 DAP-seq binding amongst the shikimate, stilbenoid, lignin and flavonoid pathways. Each gene is depicted in green, red and blue, respectively, whilst overlaps between the genes are shown with overlapping colours. The left panel shows DAP-Seq binding with a distance filter of -10kb from TSS up to +2kb after end of feature whilst the right panel shows DAP-Seq binding within -5kb from TSSs, i.e. the promoter region. The x axis shows different gene numbers (e.g. isoenzymes). A cross marks gene symbols which do not exist or are not associated to the respective pathways. An asterisk marks the manual assignment of DAP-Seq binding peaks (for the three MYBs) which were otherwise assigned by the automatic pipeline to a slightly nearer neighbouring gene.

**Figure S8:**
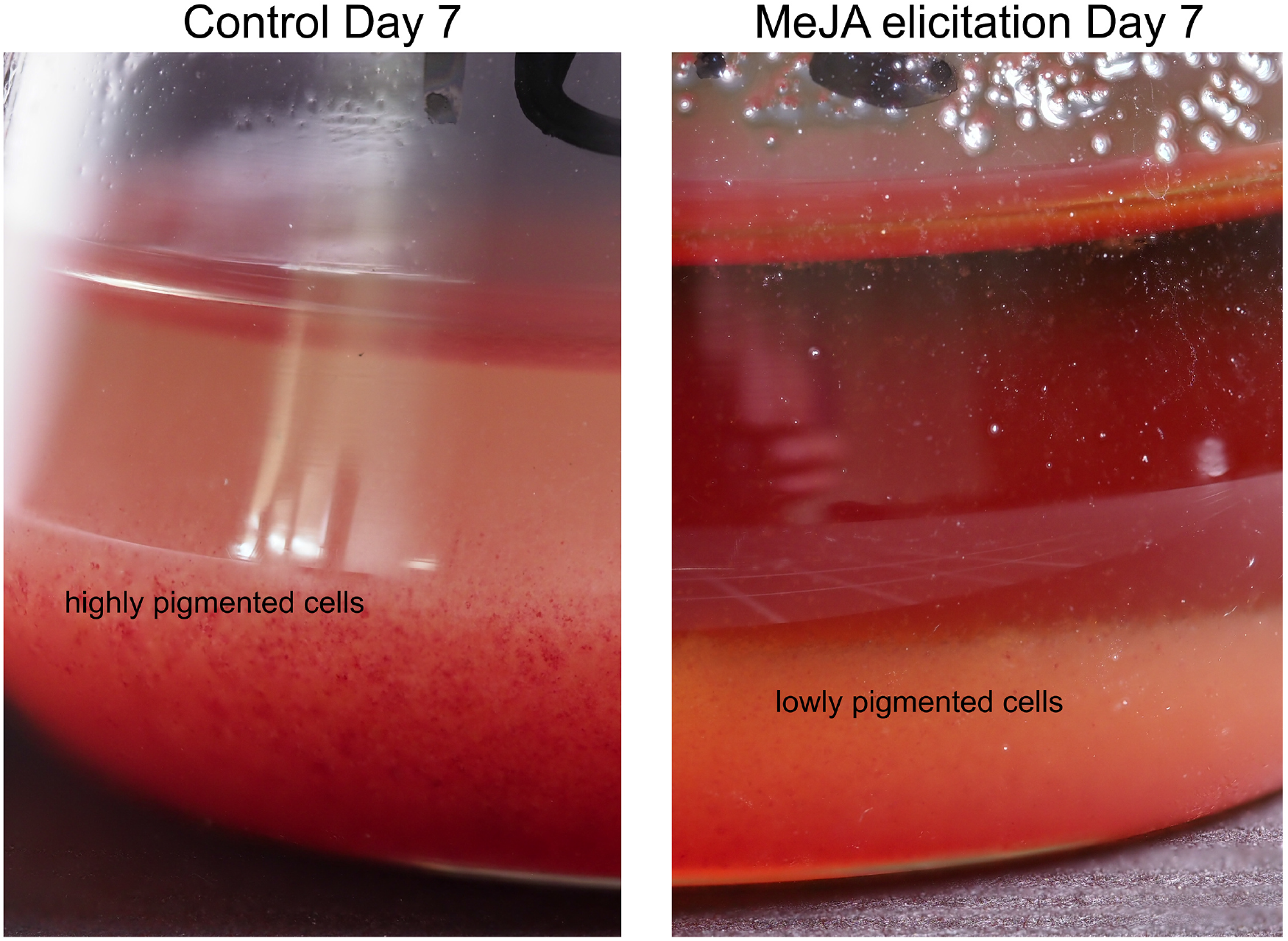
Lack of anthocyanin pigmentation in jasmonate (MeJA) and cyclodextrins (CD)-elicited grapevine cell cultures, compared to control cells at Day 7.

**Figure S9:**
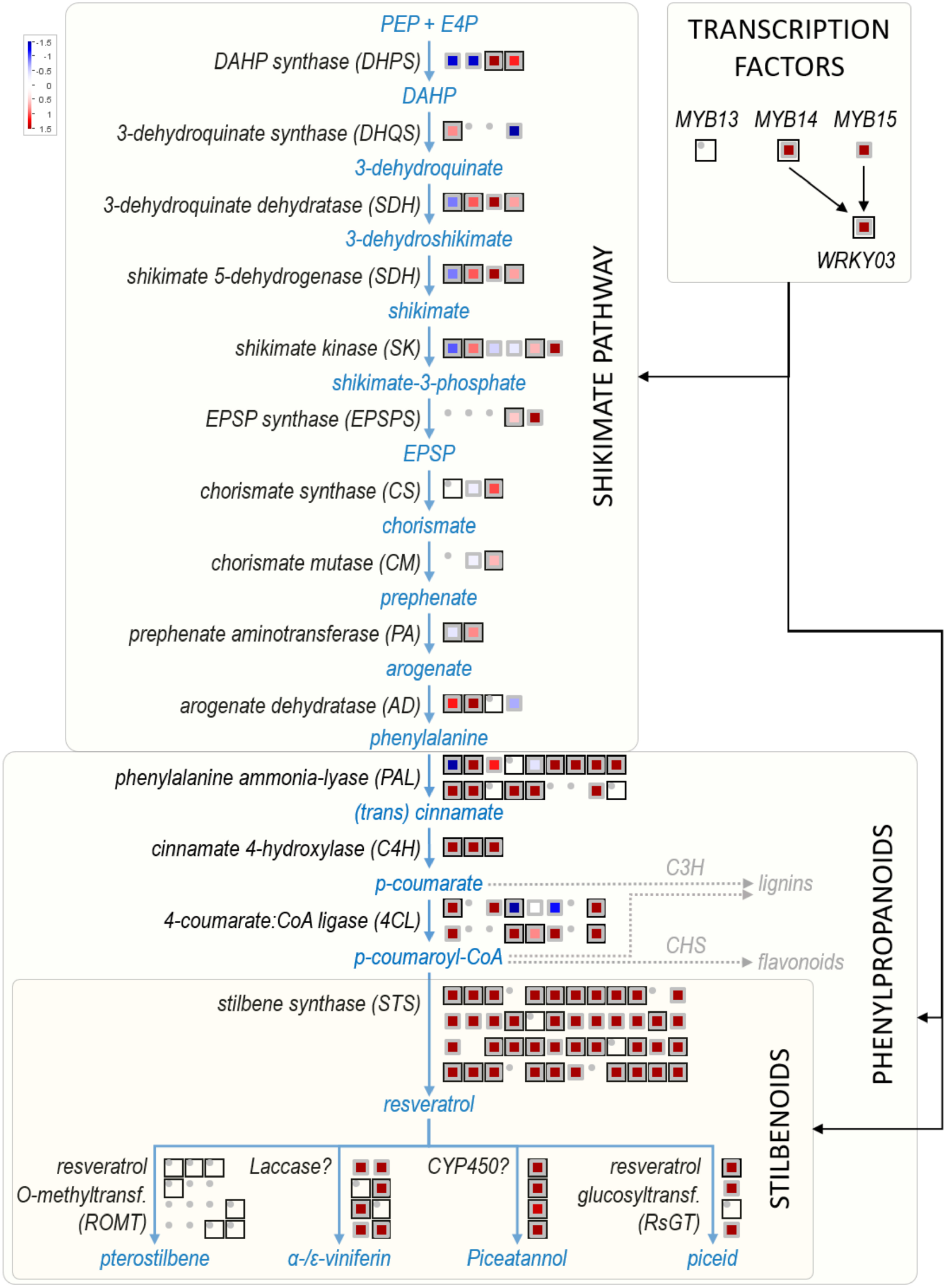
MYB13, MYB14 and MYB15 DAP-seq binding with respect to differential gene expression of shikimate, early phenylpropanoid and stilbenoid pathway genes during stage 2 noble rot infection of berries (reanalyzed from (Blanco-Ulate et al., 2015) Genes which are not bound by either VviMYB13/14/15 have grey-bordered boxes. Genes drawn with dots represent genes not passing the low-expression filter (i.e. no reads mapped). *WRKY03* is included as it has been shown to work with R2R3-MYBs to induce *STS* gene expression.

**Figure S10:**
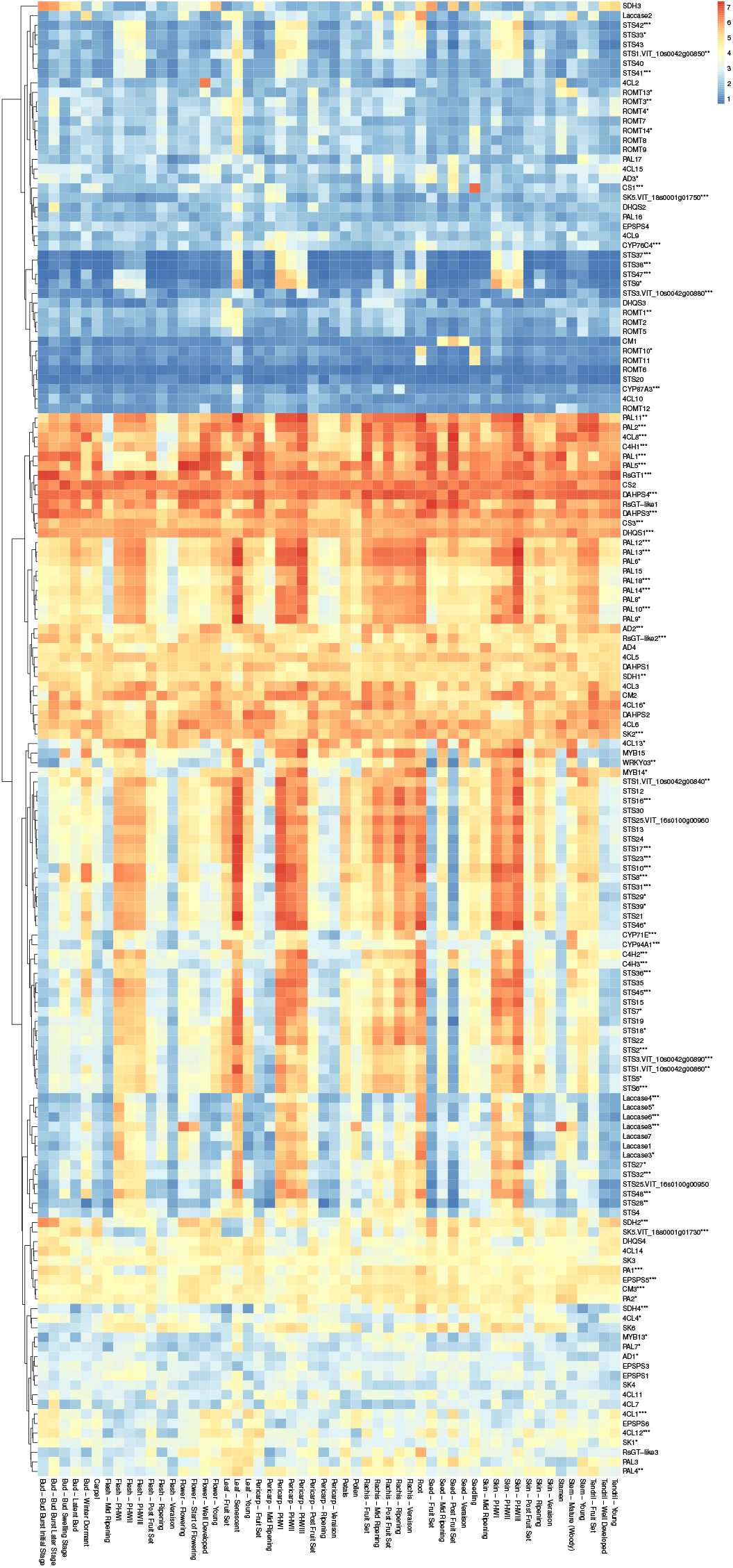
Microarray expression of shikimate, early phenylpropanoid and stilbenoid pathway genes across the *V. vinifera* cv. ‘Corvina’ expression atlas (Fasoli et al., 2012). Expression is shown as the log of (RMA + 1) across different grapevine tissues. Individual genes in the y axis are clustered according to their overall expression patterns as illustrated by the dendogram on the left. Gene symbols are based on the VCost.v3 annotation but since the microarray is based on V1 gene models, ID codes are shown where multiple V1s correspond to the same V3. DAP-Seq gene binding of MYB13/14/15 is shown by a number of asterisks at the end of gene names which correspond to the number of binding R2R3-MYBs.

**Figure S11:**
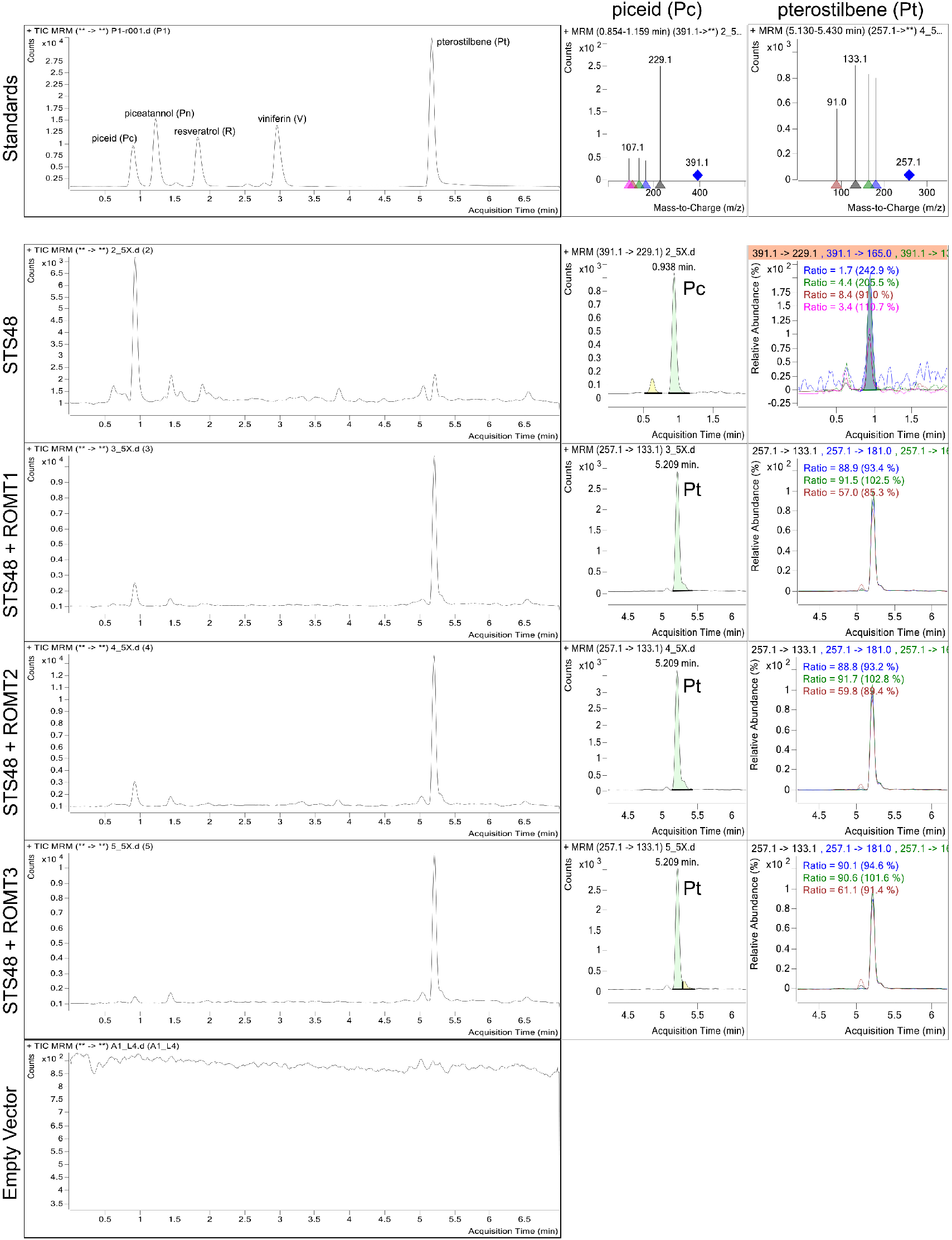
Piceid (Pc) and pterostilbene (Pt) accumulation in tobacco leaves upon overexpression of *STS48* and *ROMT1-3* genes. A comparison with the silbenoid metabolite standards shows a chromatogram peak corresponding to piced for *STS48* alone whilst pterostilbene peaks are observed for each co-transformed *ROMT1* -*3*. The different piceid and pterostilbene transitions are also shown as well as peak detection and quantification for each sample.

**Figure S12:**
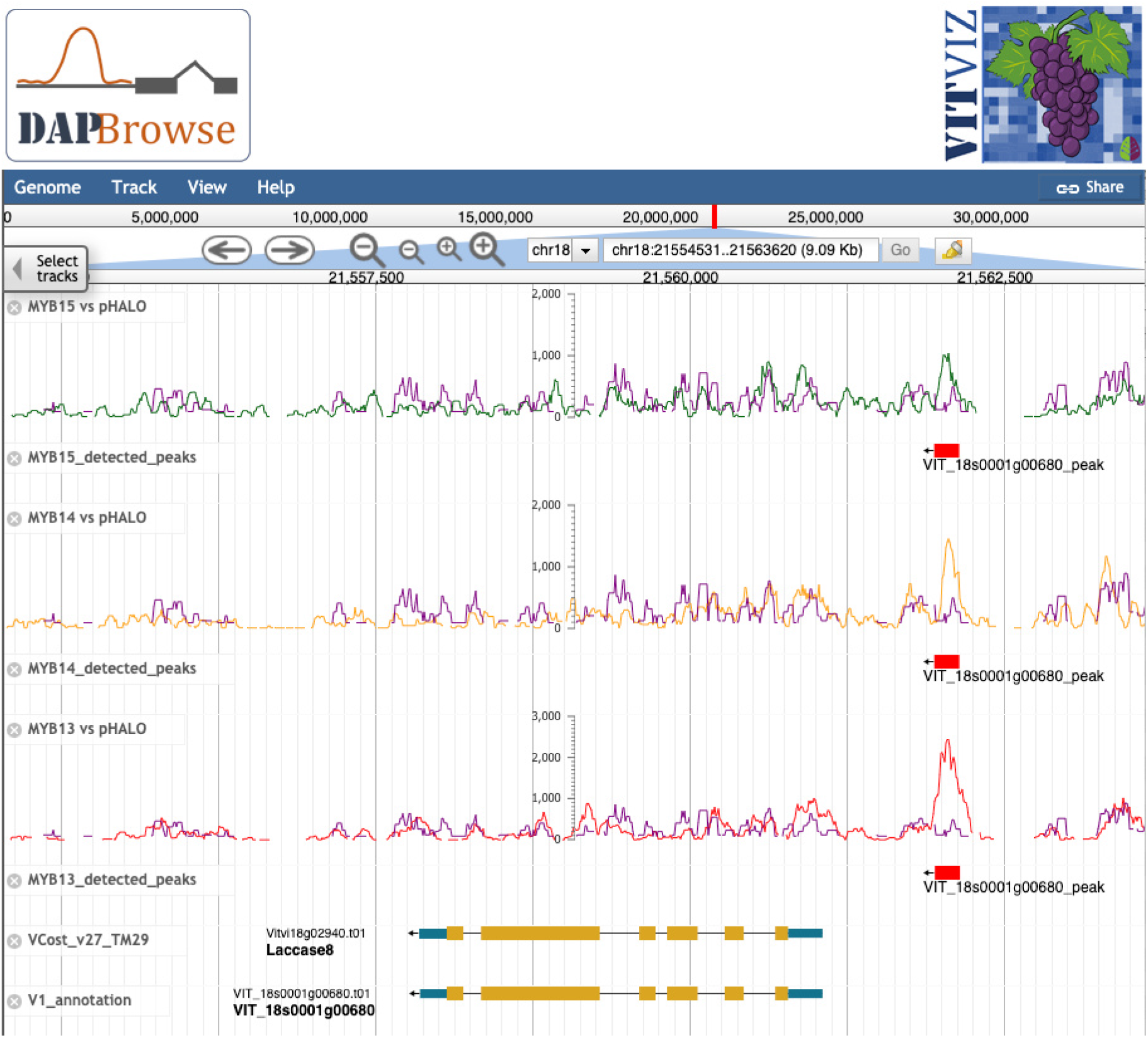
JBrowse-adaptation for the visualisation of DAP-Seq data in grapevine (DAP-Browse App, available at https://tomsbiolab.com/vitviz). JBrowse screenshot showing coverage tracks for a multireplicate DAP-Seq analysis of MYB15, MYB14 and MYB13 in the genomic regions surrounding the gene *Laccase8*. Detection of peaks and V1 and VCost.v3 annotations are included as independent tracks.

**Dataset S1** VviMYB13/14/15 DAP-Seq peaks, Gene Set Enrichment Analysis (GSEA) tables and housekeeping genes.

**Dataset S2** RNA-Seq SRA runs for network construction, *MYB14* and *MYB15* GCNs, HCTs, GSEA tables and VCost.v3 to MapMan Bin associations.

**Dataset S3** Primers, *MYB15* overexpression cluster modules, affymetrix probe-*to*-gene assignments and log_2_ fold changes, updated MapMan ontology.

## Notes

http://tomsbiolab.com/scriptsandfiles

https://tomsbiolab.com/vitviz

### Competing Interest Statement

The authors have declared no competing interest.

### Summary of Updates

This version presents additional analyses including: leaf-derived gene co-expression networks, stilbenoid elicitiation experiments in grape cell cultures and novel-enzyme characterizations.

## References

L. Almagro, P. Carbonell-Bejerano, S. Belchí-Navarro, R. Bru, J. M. Martínez-Zapater, D. Lijavetzky, and M. A. Pedreño. Dissecting the transcriptional response to elicitors in Vitis vinifera cells. PLoS ONE, 9(10):e109777, 2014. ISSN 19326203. doi: 10.1371/journal.pone.0109777.

S. Ballouz, M. Weber, P. Pavlidis, and J. Gillis. EGAD: ultra-fast functional analysis of gene networks. Bioinformatics, 33(4):612–614, 11 2016. ISSN 1367-4803. doi: 10.1093/bioinformatics/btw695. URL https://doi.org/10.1093/bioinformatics/btw695.

A. Bartlett, R. O’Malley, S.-S. Huang, M. Galli, J. Nery, A. Gallavotti, and J. Ecker. Mapping genome-wide transcription-factor binding sites using dap-seq. Nature Protocols, 12:1659–1672, 08 2017. doi: 10.1038/nprot.2017.055.

Y. Benjamini and Y. Hochberg. Controlling the false discovery rate: A practical and powerful approach to multiple testing. Journal of the Royal Statistical Society: Series B (Methodological), 57(1):289–300, 1995. doi: https://doi.org/10.1111/j.2517-6161.1995.tb02031.x. URL https://rss.onlinelibrary.wiley.com/doi/abs/10.1111/j.2517-6161.1995.tb02031.x.

B. Blanco-Ulate, K. C. Amrine, T. S. Collins, R. M. Rivero, A. R. Vicente, A. Morales-Cruz, C. L. Doyle, Z. Ye, G. Allen, H. Heymann, S. E. Ebeler, and D. Cantu. Developmental and metabolic plasticity of white-skinned grape berries in response to botrytis cinerea during noble rot. Plant Physiology, 169(4):2422–2443, 2015. ISSN 15322548. doi: 10.1104/pp.15.00852.

R. Bru, S. Sellés, J. Casado-Vela, S. Belchí-Navarro, and M. A. Pedreño. Modified cyclodextrins are chemically defined glucan inducers of defense responses in grapevine cell cultures. Journal of Agricultural and Food Chemistry, 54(1):66–71, 2006. doi: 10.1021/jf051485j.

R. Buels, E. Yao, C. M. Diesh, R. D. Hayes, M. Munoz-Torres, G. Helt, D. M. Goodstein, C. G. Elsik, S. E. Lewis, L. Stein, and I. H. Holmes. JBrowse: A dynamic web platform for genome visualization and analysis. Genome Biology, 17(1):1–12, 2016. ISSN 1474760X. doi: 10.1186/s13059-016-0924-1. URL http://dx.doi.org/10.1186/s13059-016-0924-1.

A. Canaguier, J. Grimplet, G. Di Gaspero, S. Scalabrin, E. Duchêne, N. Choisne, N. Mohellibi, C. Guichard, S. Rombauts, I. Le Clainche, A. Bérard, A. Chauveau, R. Bounon, C. Rustenholz, M. Morgante, M.-C. Le Paslier, D. Brunel, and A.-F. Adam-Blondon. A new version of the grapevine reference genome assembly (12x.v2) and of its annotation (vcost.v3). Genomics Data, 14:56 – 62, 2017. ISSN 2213-5960. doi: https://doi.org/10.1016/j.gdata.2017.09.002. URL http://www.sciencedirect.com/science/article/pii/S2213596017301459.

P. Carbonell-Bejerano, V. Rodŕıguez, C. Royo, S. Herńaiz, L. C. Moro-González, M. Torres-Viñals, and J. M. Martínez-Zapater. Circadian oscillatory transcriptional programs in grapevine ripening fruits. BMC Plant Biology, 14(1):1–15, 2014. ISSN 14712229. doi: 10.1186/1471-2229-14-78.

S. Chen, Y. Zhou, Y. Chen, and J. Gu. fastp: an ultra-fast all-in-one FASTQ preprocessor. Bioinformatics, 34(17):i884–i890, 09 2018. ISSN 1367-4803. doi: 10.1093/bioinformatics/bty560. URL https://doi.org/10.1093/bioinformatics/bty560.

Y. Chen, C. Zhangliang, J. Kang, D. Kang, H. Gu, and G. Qin. Atmyb14 regulates cold tolerance in arabidopsis. Plant Molecular Biology Reporter, 31, 02 2013. doi: 10.1007/s11105-012-0481-z.

C. S. Chin, P. Peluso, F. J. Sedlazeck, M. Nattestad, G. T. Concepcion, A. Clum, C. Dunn, R. O’Malley, R. Figueroa-Balderas, A. Morales-Cruz, G. R. Cramer, M. Delledonne, C. Luo, J. R. Ecker, D. Cantu, D. R. Rank, and M. C. Schatz. Phased diploid genome assembly with single-molecule real-time sequencing. Nature Methods, 13(12):1050–1054, 2016. ISSN 15487105. doi: 10.1038/nmeth.4035. URL http://dx.doi.org/10.1038/nmeth.4035.

A. Dobin, C. A. Davis, F. Schlesinger, J. Drenkow, C. Zaleski, S. Jha, P. Batut, M. Chaisson, and T. R. Gingeras. STAR: ultrafast universal RNA-seq aligner. Bioinformatics, 29 (1):15–21, 10 2012. ISSN 1367-4803. doi: 10.1093/bioinformatics/bts635. URL https://doi.org/10.1093/bioinformatics/bts635.

A. S. Dubrovina and K. V. Kiselev. Regulation of stilbene biosynthesis in plants. Planta, 246(4): 597–623, 2017. ISSN 14322048. doi: 10.1007/s00425-017-2730-8.

M. Fasoli, S. Dal Santo, S. Zenoni, G. B. Tornielli, L. Farina, A. Zamboni, A. Porceddu, L. Venturini, M. Bicego, V. Murino, A. Ferrarini, M. Delledonne, and M. Pezzotti. The grapevine expression atlas reveals a deep transcriptome shift driving the entire plant into a maturation program. Plant Cell, 24(9):3489–3505, 2012. ISSN 10404651. doi: 10.1105/tpc.112.100230.

Y. Guo, S. Mahony, and D. K. Gifford. High resolution genome wide binding event finding and motif discovery reveals transcription factor spatial binding constraints. PLOS Computational Biology, 8(8):1–14, 08 2012. doi: 10.1371/journal.pcbi.1002638. URL https://doi.org/10.1371/journal.pcbi.1002638.

D. Hall and V. De Luca. Mesocarp localization of a bi-functional resveratrol/hydroxycinnamic acid glucosyltransferase of Concord grape (Vitis labrusca). Plant Journal, 49(4):579–591, 2007. ISSN 09607412. doi: 10.1111/j.1365-313X.2006.02987.x.

L. Huang, X. Yin, X. Sun, J. Yang, M. Rahman, Z. Chen, and X. Wang. Expression of a grape vqsts36-increased resistance to powdery mildew and osmotic stress in arabidopsis but enhanced susceptibility to botrytis cinerea in arabidopsis and tomato. International Journal of Molecular Sciences, 19:2985, 09 2018. doi: 10.3390/ijms19102985.

J. Höll, A. Vannozzi, S. Czemmel, C. D’Onofrio, A. Walker, T. Rausch, M. Lucchin, P. Boss, I. Dry, and J. Bogs. The r2r3-myb transcription factors myb14 and myb15 regulate stilbene biosynthesis in vitis vinifera. The Plant cell, 25, 10 2013. doi: 10.1105/tpc.113.117127.

O. Jaillon, J.-M. Aury, B. Noel, A. Policriti, C. Clepet, A. Casagrande, N. Choisne, S. Aubourg, N. Vitulo, C. Jubin, A. Vezzi, F. Legeai, P. Hugueney, C. Dasilva, D. Horner, E. Mica, D. Jublot, J. Poulain, C. Bruyère, A. Billault, B. Segurens, M. Gouyvenoux, E. Ugarte, F. Cattonaro, V. Anthouard, V. Vico, C. Del Fabbro, M. Alaux, G. Di Gaspero, V. Dumas, N. Felice, S. Paillard, I. Juman, M. Moroldo, S. Scalabrin, A. Canaguier, I. Le Clainche, G. Malacrida, E. Durand, G. Pesole, V. Laucou, P. Chatelet, D. Merdinoglu, M. Delledonne, M. Pezzotti, A. Lecharny, C. Scarpelli, F. Artiguenave, M. E. Pè, G. Valle, M. Morgante, M. Caboche, A.-F. Adam-Blondon, J. Weissenbach, F. Quètier, P. Wincker, and French-Italian Public Consortium for Grapevine Genome Characterization. The grapevine genome sequence suggests ancestral hexaploidization in major angiosperm phyla. Nature, 449(7161):463—467, September 2007. ISSN 0028-0836. doi: 10.1038/nature06148. URL https://doi.org/10.1038/nature06148.

C. Jiang, D. Wang, J. Zhang, Y. Xu, C. Zhang, J. Zhang, X. Wang, and Y. Wang. VqMYB154 promotes polygene expression and enhances resistance to pathogens in Chinese wild grapevine. Horticulture Research, 8(1):1–17, 2021. ISSN 20527276. doi: 10.1038/s41438-021-00585-0. URL http://dx.doi.org/10.1038/s41438-021-00585-0.

Z. Kelemen, A. Sebastian, W. Xu, D. Grain, F. Salsac, A. Avon, N. Berger, J. Tran, B. Dubreucq, C. Lurin, L. Lepiniec, B. Contreras-Moreira, and C. Dubos. Analysis of the dna-binding activities of the arabidopsis r2r3-myb transcription factor family by one-hybrid experiments in yeast. PLOS ONE, 10(10):1–22, 10 2015. doi: 10.1371/journal.pone.0141044. URL https://doi.org/10.1371/journal.pone.0141044.

P. Kenrick and P. Crane. The origin and early evolution of plants on land. Nature, 389:33–39, 09 1997. doi: 10.1038/37918.

S. H. Kim, P. Y. Lam, M.-H. Lee, H. S. Jeon, Y. Tobimatsu, and O. K. Park. The arabidopsis r2r3 myb transcription factor myb15 is a key regulator of lignin biosynthesis in effector-triggered immunity. Frontiers in Plant Science, 11:1456, 2020. ISSN 1664-462X. doi: 10.3389/fpls.2020.583153. URL https://www.frontiersin.org/article/10.3389/fpls.2020.583153.

L. Kolberg, U. Raudvere, I. Kuzmin, J. Vilo, and H. Peterson. gprofiler2 – an R package for gene list functional enrichment analysis and namespace conversion toolset g:Profiler. F1000Research, 9:709, 2020. doi: 10.12688/f1000research.24956.2.

P. Langfelder and S. Horvath. Wgcna: an r package for weighted correlation network analysis. BMC Bioinformatics, 9:1–13, 01 2008.

B. Langmead and S. Salzberg. Langmead b, salzberg sl.. fast gapped-read alignment with bowtie 2. nat methods 9: 357-359. Nature methods, 9:357–9, 03 2012. doi: 10.1038/nmeth.1923.

F. Lecourieux, C. Kappel, P. Pieri, J. Charon, J. Pillet, G. Hilbert, C. Renaud, E. Gomès, S. Delrot, and D. Lecourieux. Dissecting the biochemical and transcriptomic effects of a locally applied heat treatment on developing Cabernet Sauvignon grape berries. Frontiers in Plant Science, 8 (JANUARY), 2017. ISSN 1664462X. doi: 10.3389/fpls.2017.00053.

Y. Liao, G. K. Smyth, and W. Shi. featureCounts: an efficient general purpose program for assigning sequence reads to genomic features. Bioinformatics, 30(7):923–930, 11 2013. ISSN 1367-4803. doi: 10.1093/bioinformatics/btt656. URL https://doi.org/10.1093/bioinformatics/btt656.

P. Machanick and T. Bailey. Meme-chip: motif analysis of large dna datasets. *Bioinformatics (Oxford*,. England), 27:1696–7, 06 2011. doi: 10.1093/bioinformatics/btr189.

A. Martínez-Márquez, J. A. Morante-Carriel, K. Ramírez-Estrada, R. M. Cusidó, J. Palazon, and R. Bru-Martínez. Production of highly bioactive resveratrol analogues pterostilbene and piceatannol in metabolically engineered grapevine cell cultures. Plant Biotechnology Journal, 14 (9):1813–1825, 2016. ISSN 14677652. doi: 10.1111/pbi.12539.

P. R. Merz, T. Moser, J. Höll, A. Kortekamp, G. Buchholz, E. Zyprian, and J. Bogs. The transcription factor VvWRKY33 is involved in the regulation of grapevine (Vitis vinifera) defense against the oomycete pathogen Plasmopara viticola. Physiologia Plantarum, 153(3):365–380, 2015. ISSN 13993054. doi: 10.1111/ppl.12251.

M. Mutwil, S. Klie, T. Tohge, F. M. Giorgi, O. Wilkins, M. M. Campbell, A. R. Fernie, B. Usadel, Z. Nikoloski, and S. Persson. PlaNet: Combined Sequence and Expression Comparisons across Plant Networks Derived from Seven Species . The Plant Cell, 23(3):895–910, 03 2011. ISSN 1040-4651. doi: 10.1105/tpc.111.083667. URL https://doi.org/10.1105/tpc.111.083667.

M. Nakata and M. Ohme-Takagi. Quantification of Anthocyanin Content. Bio-protocol, 4(9):e1098, 2014. doi: 10.21769/BioProtoc.1098.

R. O’Malley, S. shan Carol Huang, L. Song, M. Lewsey, A. Bartlett, J. Nery, M. Galli, A. Gallavotti, and J. Ecker. Cistrome and epicistrome features shape the regulatory dna landscape. Cell, 165 (5):1280 – 1292, 2016. ISSN 0092-8674. doi: https://doi.org/10.1016/j.cell.2016.04.038. URL http://www.sciencedirect.com/science/article/pii/S0092867416304810.

R. Parage, R. Tavares, S. Réty, R. Baltenweck-Guyot, A. Poutaraud, L. Renault, D. Heintz, Lugan, G. A. Marais, S. Aubourg, and P. Hugueney. Structural, functional, and evolutionary analysis of the unusually large stilbene synthase gene family in grapevine. Plant Physiology, 160(3):1407–1419, 2012. ISSN 0032-0889. doi: 10.1104/pp.112.202705. URL http://www.plantphysiol.org/content/160/3/1407.

S. Pasquereau, Z. Nehme, S. Haidar Ahmad, F. Daouad, J. Van Assche, C. Wallet, C. Schwartz, O. Rohr, S. Morot-Bizot, and G. Herbein. Resveratrol Inhibits HCoV-229E and SARS-CoV-2 Coronavirus Replication In Vitro. Viruses, 13(2):1–11, 2021. ISSN 1999-4915. doi: 10.3390/v13020354. URL http://www.ncbi.nlm.nih.gov/pubmed/33672333.

C. Qian, Z. Chen, Q. Liu, W. Mao, Y. Chen, W. Tian, Y. Liu, J. Han, X. Ouyang, and X. Huang. Coordinated transcriptional regulation by the uv-b photoreceptor and multiple transcription factors for plant uv-b responses. Molecular Plant, 13(5):777–792, 2020. ISSN 1674-2052. doi: https://doi.org/10.1016/j.molp.2020.02.015. URL https://www.sciencedirect.com/science/article/pii/S1674205220300629.

Ž. Ramšak, Š. Baebler, A. Rotter, M. Korbar, I. Mozetič, B. Usadel, and K. Gruden. GoMapMan: Integration, consolidation and visualization of plant gene annotations within the MapMan ontology. Nucleic Acids Research, 42(D1):1167–1175, 2014. ISSN 03051048. doi: 10.1093/nar/gkt1056.

M. E. Ritchie, B. Phipson, D. Wu, Y. Hu, C. W. Law, W. Shi, and G. K. Smyth. limma powers differential expression analyses for RNA-sequencing and microarray studies. Nucleic Acids Research, 43(7):e47–e47, 01 2015. ISSN 0305-1048. doi: 10.1093/nar/gkv007. URL https://doi.org/10.1093/nar/gkv007.

I. Romero, A. Fuertes, M. J. Benito, J. M. Malpica, A. Leyva, and J. Paz-Ares. More than 80R2R3-MYB regulatory genes in the genome of Arabidopsis thaliana. Plant Journal, 14(3):273–284, 1998. ISSN 09607412. doi: 10.1046/j.1365-313X.1998.00113.x.

M. Santos-Rosa, A. Poutaraud, D. Merdinoglu, and P. Mestre. Development of a transient expression system in grapevine via agro-infiltration. Plant Cell Reports, 27(6):1053–1063, 2008. ISSN 07217714. doi: 10.1007/s00299-008-0531-z.

L. Schmidlin, A. Poutaraud, P. Claudel, P. Mestre, E. Prado, M. Santos-Rosa, S. Wiedemann-Merdinoglu, F. Karst, D. Merdinoglu, and P. Hugueney. A stress-inducible resveratrol O-methyltransferase involved in the biosynthesis of pterostilbene in grapevine. Plant Physiology, 148(3):1630–1639, 2008. ISSN 00320889. doi: 10.1104/pp.108.126003.

R. Schwacke, G. Y. Ponce-Soto, K. Krause, A. M. Bolger, B. Arsova, A. Hallab, K. Gruden, M. Stitt, M. E. Bolger, and B. Usadel. MapMan4: A Refined Protein Classification and Annotation Framework Applicable to Multi-Omics Data Analysis. Molecular Plant, 12(6):879–892, 2019. ISSN 17529867. doi: 10.1016/j.molp.2019.01.003.

A. Shalit-Kaneh, R. W. Kumimoto, V. Filkov, and S. L. Harmer. Multiple feedback loops of the Arabidopsis circadian clock provide rhythmic robustness across environmental conditions. Proceedings of the National Academy of Sciences of the United States of America, 115(27):7147–7152, 2018. ISSN 10916490. doi: 10.1073/pnas.1805524115.

O. Thimm, O. Bläsing, Y. Gibon, A. Nagel, S. Meyer, P. Krüger, J. Selbig, L. A. Müller, S. Y. Rhee, and M. Stitt. MAPMAN: A user-driven tool to display genomics data sets onto diagrams of metabolic pathways and other biological processes. The Plant Journal, 37(6):914–939, 2004. ISSN 09607412. doi: 10.1111/j.1365-313X.2004.02016.x.

A. Vannozzi, I. Dry, M. Fasoli, S. Zenoni, and M. Lucchin. Genome-wide analysis of the grapevine stilbene synthase multigenic family: Genomic organization and expression profiles upon biotic and abiotic stresses. BMC plant biology, 12:130, 08 2012. doi: 10.1186/1471-2229-12-130.

A. Vannozzi, D. C. J. Wong, J. Höll, I. Hmmam, J. T. Matus, J. Bogs, T. Ziegler, I. Dry, G. Barcaccia, and M. Lucchin. Combinatorial Regulation of Stilbene Synthase Genes by WRKY and MYB Transcription Factors in Grapevine (Vitis vinifera L.). Plant and Cell Physiology, 59(5):1043–1059, 2018. ISSN 14719053. doi: 10.1093/pcp/pcy045.

W. Verleyen, S. Ballouz, and J. Gillis. Measuring the wisdom of the crowds in network-based gene function inference. *Bioinformatics (Oxford*. England), 31, 10 2014. doi: 10.1093/bioinformatics/btu715.

D. Wang, C. Jiang, R. Li, and Y. Wang. VqbZIP1 isolated from Chinese wild Vitis quinquangularis is involved in the ABA signaling pathway and regulates stilbene synthesis. Plant Science, 287(July):110202, 2019. ISSN 18732259. doi: 10.1016/j.plantsci.2019.110202. URL https://doi.org/10.1016/j.plantsci.2019.110202.

D. Wang, C. Jiang, W. Liu, Y. Wang, and R. Hancock. The WRKY53 transcription factor enhances stilbene synthesis and disease resistance by interacting with MYB14 and MYB15 in Chinese wild grape. Journal of Experimental Botany, 71(10):3211–3226, 2020. ISSN 14602431. doi: 10.1093/jxb/eraa097.

L. Wang and Y. Wang. Transcription factor VqERF114 regulates stilbene synthesis in Chinese wild Vitis quinquangularis by interacting with VqMYB35. Plant Cell Reports, 38(10):1347–1360, 2019. ISSN 1432203X. doi: 10.1007/s00299-019-02456-4. URL https://doi.org/10.1007/s00299-019-02456-4.

E. R. Waters. Molecular adaptation and the origin of land plants. Molecular Phylogenetics and Evolution, 29(3):456 – 463, 2003. ISSN 1055-7903. doi: https://doi.org/10.1016/j.ympev.2003.07.018. URL http://www.sciencedirect.com/science/article/pii/S1055790303003130. Plant Molecular Evolution.

C. J. Wolfe, I. S. Kohane, and A. J. Butte. Systematic survey reveals general applicability of ”guilt-by-association” within gene coexpression networks. BMC Bioinformatics, 6:1–10, 2005. ISSN 14712105. doi: 10.1186/1471-2105-6-227.

D. Wong. Network aggregation improves gene function prediction of grapevine gene co-expression networks. Plant Molecular Biology, 103(4-5):425–441, 2020. ISSN 15735028. doi: 10.1007/s11103-020-01001-2. URL https://doi.org/10.1007/s11103-020-01001-2.

D. C. J. Wong and J. T. Matus. Constructing integrated networks for identifying new secondary metabolic pathway regulators in grapevine: Recent applications and future opportunities. Frontiers in Plant Science, 8(April):1–8, 2017. ISSN 1664462X. doi: 10.3389/fpls.2017.00505.

D. C. J. Wong, R. Schlechter, A. Vannozzi, J. Höll, I. Hmmam, J. Bogs, G. B. Tornielli, S. D. Castellarin, and J. T. Matus. A systems-oriented analysis of the grapevine R2R3-MYB transcription factor family uncovers new insights into the regulation of stilbene accumulation. DNA Research, 23(5):451–466, 2016. ISSN 17561663. doi: 10.1093/dnares/dsw028.

H. Xi, L. Ma, G. Liu, N. Wang, J. Wang, L. Wang, Z. Dai, S. Li, and L. Wang. Transcriptomic analysis of grape (Vitis vinifera L.) leaves after exposure to ultraviolet C irradiation. PLoS ONE, 9(12), 2014. ISSN 19326203. doi: 10.1371/journal.pone.0113772.

W. Xu, M. Fuli, R. Li, Q. Zhou, W. Yao, Y. Jiao, C. Zhang, J. Zhang, X. Wang, Y. Xu, and Y. Wang. Vpsts29/sts2 enhances fungal tolerance in grapevine through a positive feedback loop. Plant, Cell & Environment, 42, 07 2019. doi: 10.1111/pce.13600.

X. Yin, S. D. Singer, H. Qiao, Y. Liu, C. Jiao, H. Wang, Z. Li, Z. Fei, Y. Wang, C. Fan, and X. Wang. Insights into the mechanisms underlying ultraviolet-c induced resveratrol metabolism in grapevine (V. amurensis rupr.) cv. “Tonghua-3”. Frontiers in Plant Science, 7(APR2016):1–16, 2016. ISSN 1664462X. doi: 10.3389/fpls.2016.00503.

L. Zhu, C. Gazin, N. Lawson, H. Pagès, S. Lin, D. Lapointe, and M. Green. Chippeakanno: A bioconductor package to annotate chip-seq and chip-chip data. BMC bioinformatics, 11:237, 05 2010. doi: 10.1186/1471-2105-11-237.

